# Aging is associated with a systemic length-driven transcriptome imbalance

**DOI:** 10.1101/691154

**Authors:** Thomas Stoeger, Rogan A. Grant, Alexandra C. McQuattie-Pimentel, Kishore Anekalla, Sophia S. Liu, Heliodoro Tejedor-Navarro, Benjamin D. Singer, Hiam Abdala-Valencia, Michael Schwake, Marie-Pier Tetreault, Harris Perlman, William E Balch, Navdeep Chandel, Karen Ridge, Jacob I. Sznajder, Richard I. Morimoto, Alexander V. Misharin, G.R. Scott Budinger, Luis A. Nunes Amaral

**Affiliations:** Department of Chemical and Biological Engineering, Northwestern University; Northwestern Institute on Complex Systems, Northwestern University; Center for Genetic Medicine, Northwestern University; Department of Molecular Biosciences, Northwestern University; Division of Pulmonary and Critical Care Medicine, Northwestern University; Department of Biochemistry and Molecular Genetics, Northwestern University; Department of Neurology, Northwestern University; Division of Gastroenterology and Hepatology, Northwestern University; Division of Rheumatology, Northwestern University; The Scripps Research Institute; Department of Physics and Astronomy, Northwestern University

## Abstract

Aging manifests itself through a decline in organismal homeostasis and a multitude of cellular and physiological functions^1^. Efforts to identify a common basis for vertebrate aging face many challenges; for example, while there have been documented changes in the expression of many hundreds of mRNAs, the results across tissues and species have been inconsistent^2^. We therefore analyzed age-resolved transcriptomic data from 17 mouse organs and 51 human organs using unsupervised machine learning^3–5^ to identify the architectural and regulatory characteristics most informative on the differential expression of genes with age. We report a hitherto unknown phenomenon, a systemic age-dependent length-driven transcriptome imbalance that for older organisms disrupts the homeostatic balance between short and long transcript molecules for mice, rats, killifishes, and humans. We also demonstrate that in a mouse model of healthy aging, length-driven transcriptome imbalance correlates with changes in expression of *splicing factor proline and glutamine rich* (*Sfpq*), which regulates transcriptional elongation according to gene length^6^. Furthermore, we demonstrate that length-driven transcriptome imbalance can be triggered by environmental hazards and pathogens. Our findings reinforce the picture of aging as a systemic homeostasis breakdown and suggest a promising explanation for why diverse insults affect multiple age-dependent phenotypes in a similar manner.

## Main text

The transcriptome responds rapidly, selectively, strongly, and reproducibly to a wide variety of molecular and physiological insults experienced by an organism^7^. While the transcripts of thousands of genes have been reported to change with age^2^, the magnitude by which most transcripts change is small in comparison with classical examples of gene regulation^2,8^ and there is little consensus among different studies. We hence hypothesize that aging is associated with a hitherto uncharacterized process that affects the transcriptome in a systemic manner. We predict that such a process could integrate heterogenous, and molecularly distinctive, environmental insults to promote phenotypic manifestations of aging^1^.

We use an unsupervised machine learning approach^3–5^ to identify the sources of age-dependent changes in the transcriptome. To this end, we measure and survey the transcriptome of 17 mouse organs from 6 biological replicates at 5 different ages from 4 to 24 months raised under standardized conditions (Fig. 1A). We consider information on the structural architecture of individual genes and transcripts, and knowledge on the binding of regulatory molecules such as transcription factors and microRNAs (miRNAs) (Fig. 1B). We define age-dependent fold-changes as the log_2_-transformed ratio of transcripts of one gene at a given age relative to the transcripts of that gene in the organs of 4-month-old mice. As expected for models capturing most measurable changes in transcript abundance, the predicted fold-changes (Fig. S1) match changes empirically observed between distinct replicate cohorts of mice (Figs. S2 and S3).

**Fig. 1.**
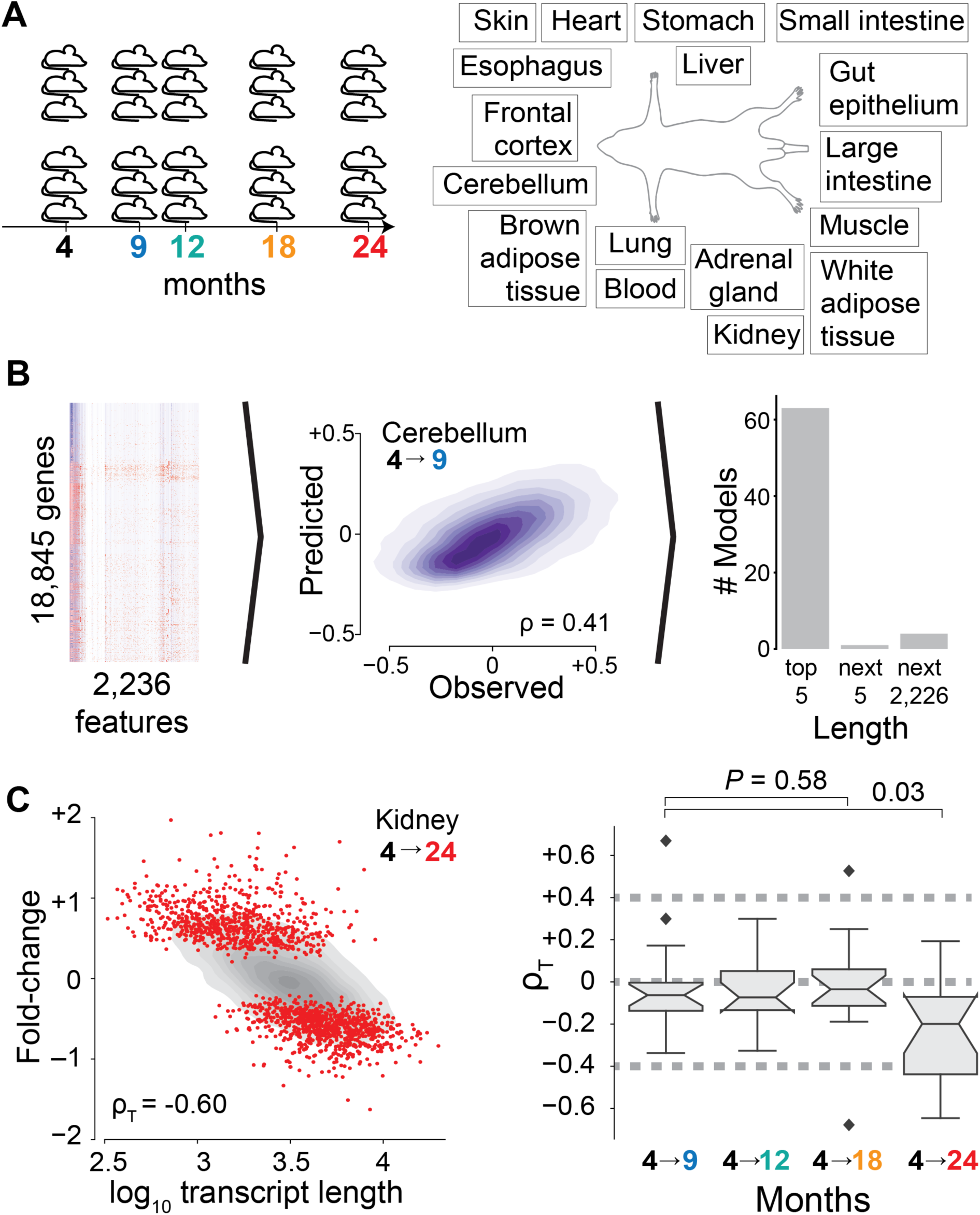
Discovery of length-driven transcriptome imbalance in aging. (**A**) At 4, 9, 12, 18, and 24 months of age, mice were sacrificed in two cohorts of three mice each and assayed by RNA sequencing for the listed organs. (**B**) A machine learning (ML) model was developed to predict fold-change of transcripts between samples from two ages for a given tissue using 2,236 features corresponding to known gene-specific regulators (transcription factors, miRNAs) and structural characteristics of genes and transcripts. A high-performance example of ML model is shown (middle panel). Analysis of features with greatest impact on performance of ML model (right panel) shows that length (median transcript length, gene length, or median length of coding sequence) consistently ranks among the most important features across all tissue- and age-specific models. (**C**) Dependence of fold-change observed between kidney samples from 4- and 24-month-old mice on median transcript length. Transcriptome imbalance (ρ_T_) was defined at the Spearman correlation of the data. Grey shows kernel density estimate of all genes whereas red dots highlight genes identified by gene-specific differential expression. Right panel shows transcriptome imbalance (ρ_T_) for the 17 organs of our study. *P* values were estimated by two-sided Mann-Whitney *U* tests.

Further supporting the sensitivity of our machine learning approach, transcriptome-wide predictions reach statistical significance in 9-month-old organs (Fig. S4) for which complementary gene-specific differential gene expression analyses had not yet identified any age-regulated gene (Fig. 1B, Figs. S5 and S6). This demonstrates that architectural and regulatory features of genes inform on age-dependent changes of the transcriptome across multiple organs.

To identify whether there are universal architectural or regulatory features informative on age-dependent changes, we systematically analyze feature importance across models. The most informative feature to those models is the median length of mature transcript molecules (Fig. 1B, Table S1), which is closely followed by the number of transcription factors, the length of the gene, and the median length of the coding sequence (see Fig. S7 for additional details). We conclude that during aging, transcript length is the most informative feature.

To determine whether transcript length could directly associate with age-dependent changes of the transcriptome, or whether transcript length solely contributes to our models through combinatorial interactions with additional architectural or regulatory features, we directly compare observed fold-changes against transcript length. We find significant support (at *P* values of <10^-40^) for such a direct association for every organ (Fig. S8). For several organs, such as 24-month-old kidneys, this relation is visually apparent (Fig. 1C, Figs. S9 and S10). For the vast majority of organs, we find that expression changes and transcript length are anticorrelated, indicating a systematic upregulation of short transcripts with age and a systematic downregulation of long transcripts, which is already visible for 9-month-old animals and further increases for 18- and 24-month-old animals (Fig. 1C). To emphasize the systemic nature of the anticorrelation between transcript length and fold-change in older animals, we term this phenomenon “length-driven transcriptome imbalance.”

To determine whether length-driven transcriptome imbalance extends beyond our own experimental conditions in mice, we next inspect genes reported to be down- and upregulated with age within recent surveys of vertebrates^5,9,10^. In an independent mouse study, we can confirm a reduced expression among long transcripts that change across multiple organs, and for 2 of 4 organ-specific sets of transcripts (differentially expressed across 3-, 12-, and 29-month-old animals). Furthermore, we find a reduced expression of long transcripts in older animals for 3 of 3 killifish organs (differentially expressed between 5- and 39-week-old animals), and 10 of 11 rat organs (8 organs between 2- and 6-week-old animals, 3 organs between 6- and 21-week-old animals, and 2 organs between 21- and 104-week-old animals) (Fig. 2A, Fig. S11).

**Fig. 2.**
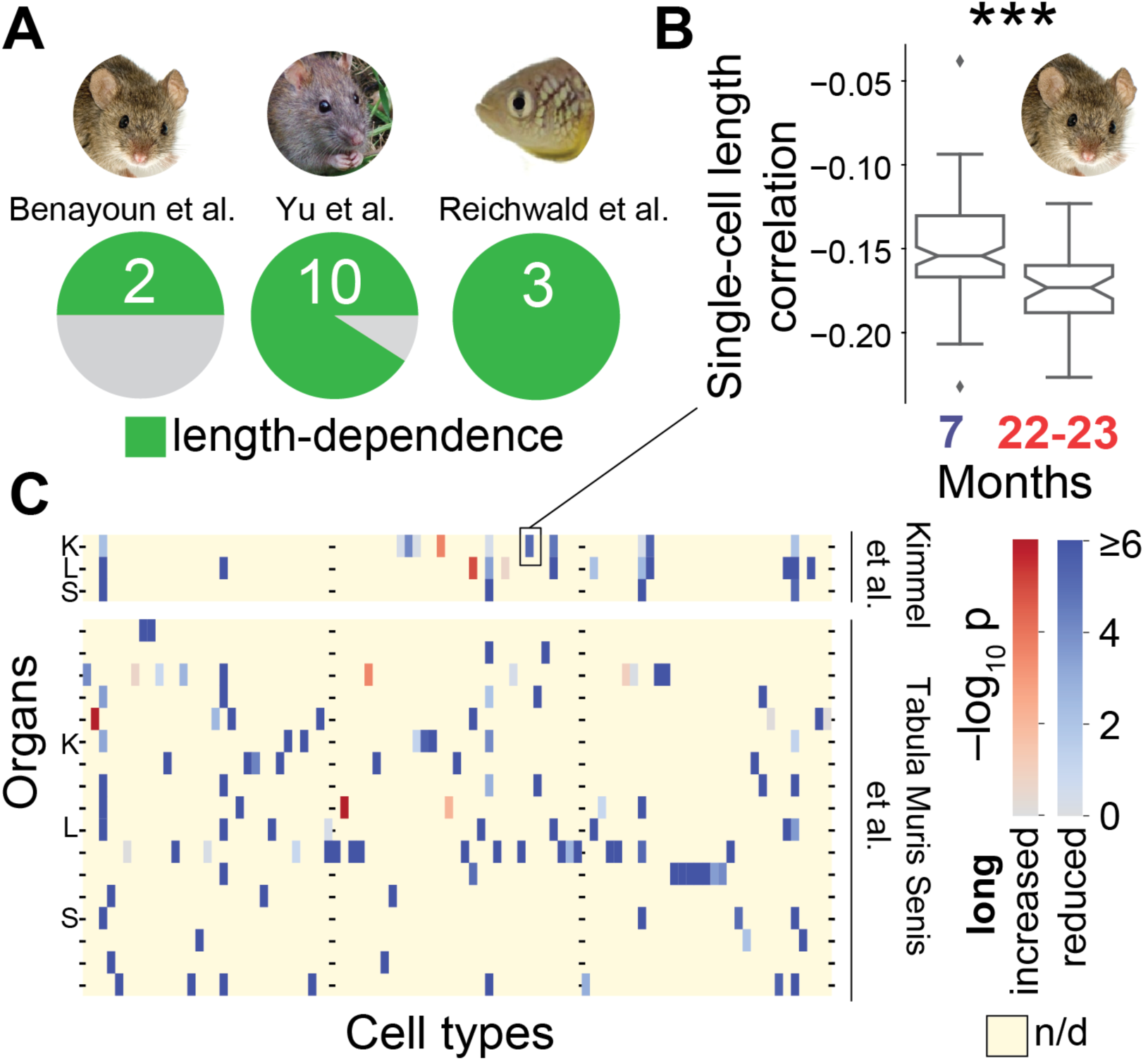
Length-driven transcriptome imbalance in other vertebrates. (**A**) Genes reported as significantly up- and downregulated with age significantly differ in the median length of their transcripts in 2 of 4 mouse organs, 10 of 11 rat organs, and 3 of 3 killifish organs (*P* < 0.05; two-sided Mann-Whitney *U* test). (**B**) Increased imbalance in old kidney mesangial cells. Single-cell length correlations are defined as the spearman correlations between transcript length and transcript levels. \*\*\**P* < 0.001 in two-sided Mann-Whitney *U* test. (**C**) Significance of shifted single-cell length correlations between older and younger animals for data by Kimmel et al.^11^ and Tabula Muris Senis et al.^12^; Red indicates that median single-cell length correlations are lower in older animals and blue indicates that median single-cell length correlations are higher in older animals. Yellow indicates absence of data (e.g.: if cell type absent or not detected); K is kidney, L is lung, S is spleen.

To resolve, whether the length-driven imbalance observed among the bulk-transcriptomes of entire organs reflected upon a change of cellular composition or a molecular process occurring in a subset of cell types, we next reanalyze the data of a recent preprint which measured transcriptomes for single cells of three organs of 7-month-old and 22-23-month-old mice^11^. For every single cell we correlate transcript lengths against the level of expression. In strong support of systemic changes, we observe a significant reduction of transcripts from long genes among single cells of 6 of 11 cell types of the kidney – including cell types with organ-specific roles such as mesangial cells (Fig. 2B, Fig. S12) –, 10 of 12 cell types of the lung, and 4 of 4 cell types of the spleen. When reanalyzing the data provided for 17 mouse organs by another recent preprint by the Tabula Muris Senis consortium^12^, we find 93 of 111 cell types to have a significant reduction of long transcripts in 24-month-old animals relative to 3-month-old animals (Fig. 2C, Fig. S13). We conclude that an imbalance of transcripts occurs in most cell types of mice, and – paralleling our earlier findings on entire organs – mainly disfavors long genes in aged individuals.

To determine, whether length-driven transcriptome imbalance also occurred during human aging, we next reanalyze transcriptomes collected by the GTEx consortium^13^ generated from human tissues at the time of their death (Fig. 3A). Supporting our initial findings in mice, machine learning models trained on those transcriptomes recover gene length as the most informative feature (Fig. 3B). Also informative were the guanine-cytosine (GC) content of transcripts, the number of different transcription factors bound to the transcriptional start site, and the length of the transcript molecules (Figs. S14 and S15, Table S2). Further matching our findings on mice, length-driven transcriptome imbalance is already apparent for the majority of organs among middle-aged donors (40–59 years) in comparison with young donors (20–39 years). In contrast to our own experiments, GTEx sampled several regions of individual organs, revealing that, among humans, transcriptome imbalance is most pervasive in the brain (Fig. 3D) both among female and male donors (Fig. S16). We conclude that transcriptome imbalance occurs in multiple vertebrates, including humans, and affects individual organs to a varying extent.

**Fig. 3.**
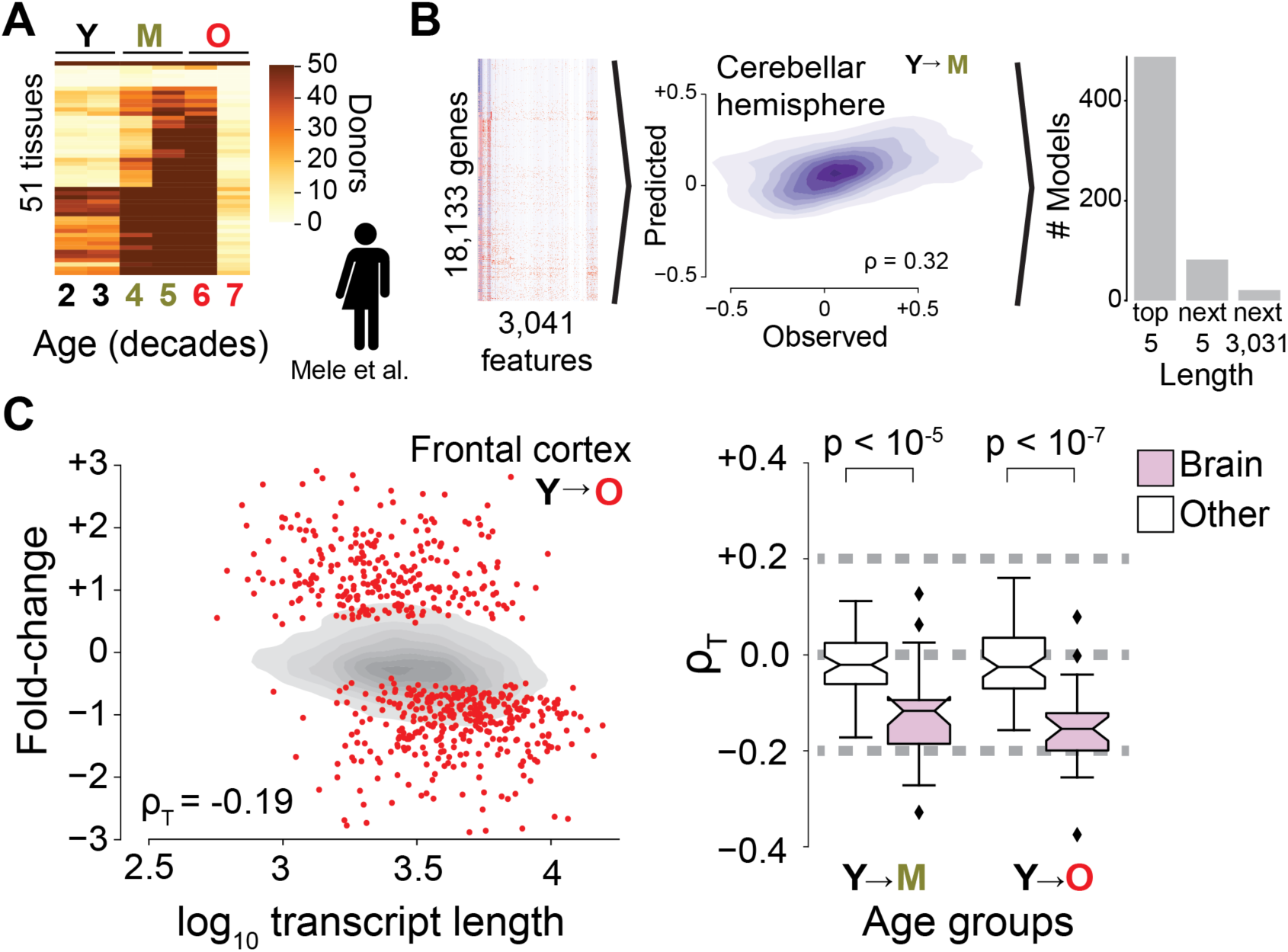
Length-driven transcriptome imbalance in humans. (**A**) Number of samples archived by the GTEx consortium for individual tissues as function of donor age. “Y” marks donors aged 20–39 years; “M” marks donors aged 40–59 years; “O” marks donors aged 60–79 years. (**B**) Same as Fig. 1B, but for human regulatory elements, and transcriptomes measured by the GTEx consortium. (**C**) Transcriptome imbalance in human GTEx. Pink box plots are for tissues from different brain regions. *P* values were estimated by two-sided Mann-Whitney *U* tests

We hypothesize that transcriptome imbalance may primarily challenge or promote cellular processes important to aging. Since length-driven transcriptome imbalance most strongly affects short and long transcripts (Fig. 4A, Fig. S17), we focus on them. Consistent with current knowledge, the mapping of known longevity mutants recovered from model organisms onto human and mouse genes yields a finding that the shortest transcripts are significantly depleted from genes beneficial to longevity and significantly enriched for deleterious effects, whereas long transcripts are opposingly enriched for beneficial effects and opposingly depleted from deleterious effects (Fig. 4B, Fig. S18). More broadly, Gene Ontology analysis for annotations enriched among transcripts of one length extreme and simultaneously depleted among transcripts of the other length extreme recover well-established facts in the literature concerning molecular, cellular, and physiological processes associated with aging (Tables S3– S6). Short genes are enriched for proteostasis, mitochondrial function, telomere maintenance, chromatin organization, and immune function^14,15^. Long genes are enriched for transcriptional regulation^16^, developmental processes^17^, ATP binding^18^, cytoskeletal structure, and synaptic activity^19^ (Fig. 4C). Collectively, these findings demonstrate a remarkably high overlap between the functions encoded by the shortest and longest transcripts and the biological hallmarks of aging^15,20^.

**Fig. 4.**
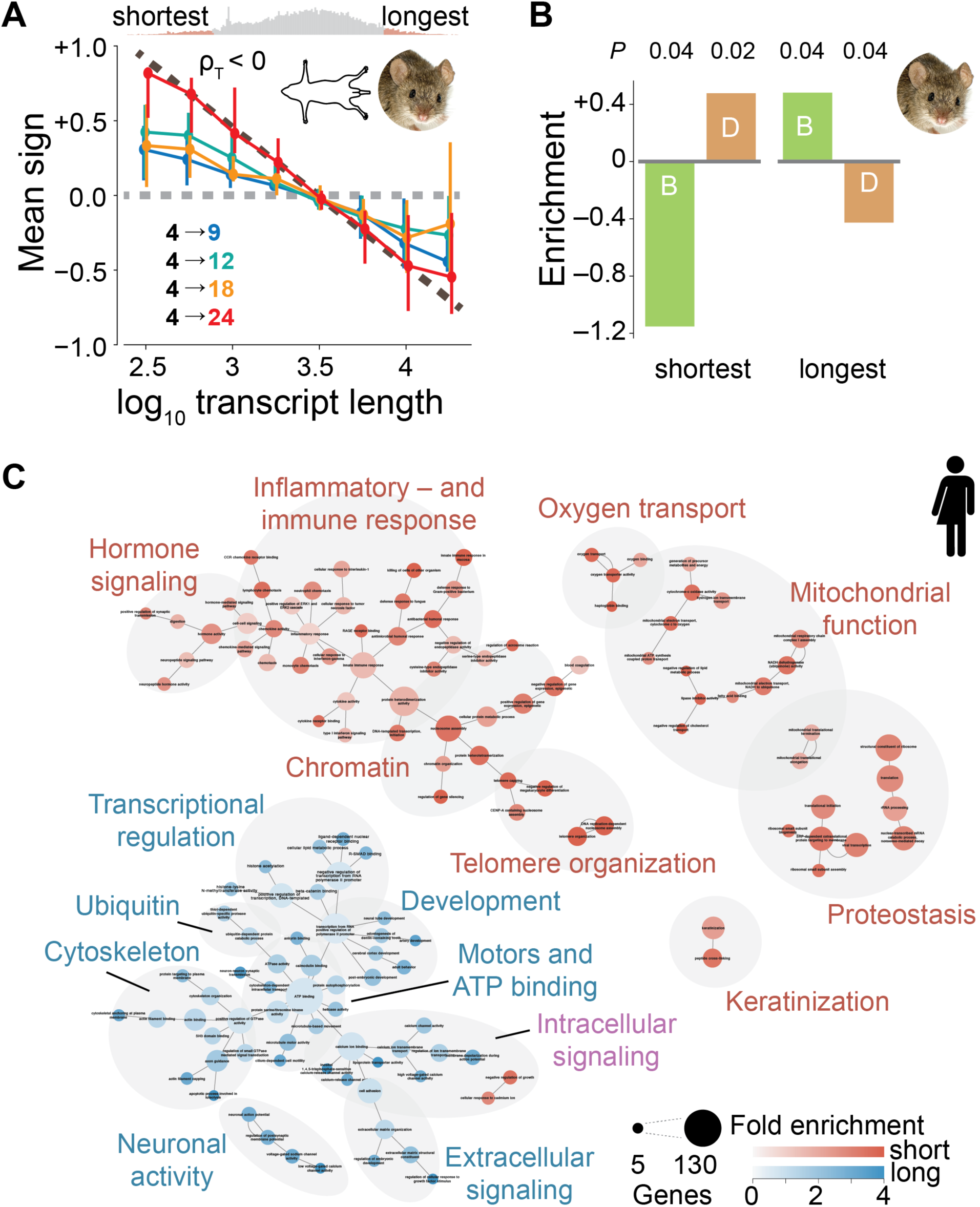
Short and long transcripts enrich for biological processes previously associated with aging. (**A**) Direction of fold-change for transcripts relative to 4-month-old mice across all organs. An average sign of +1 would indicate that all genes are upregulated, whereas an average sign of −1 would indicate that all are downregulated. Colors indicate age comparisons as in Fig. 1A. Circles show median values of fold-change across all samples and error bars represent 95% confidence intervals. Dashed brown line is linear approximation to slope seen in 24-month-old animals. Histogram shows genes with indicated transcript length. Genes with the 5% shortest and 5% longest median lengths are colored in red. Note the visible and significant negative correlation between transcript length and fold-change. (**B**) Fold enrichment for beneficial (B, green) and deleterious (D, orange) genes among the genes with the 5% shortest and 5% longest median transcript lengths in humans. Negative enrichment indicates depletion. (**C**) Human Gene Ontology analysis for annotations enriched (depleted) among genes within transcripts in the bottom (top) 5% of gene lengths. Area of circle is proportional to number of enriched genes. Edges represent highest embedding of a smaller annotation within a larger one. Red (blue) indicates genes enriched in genes with shortest (longest) transcripts (*P* < 0.05; Benjamini-Hochberg corrected Fisher’s exact test).

Next, we investigate possible origins of transcriptome imbalance. Genes whose transcript expressions correlate with transcriptome imbalance beyond the correlation expected by their own transcript length (Fig. 5A) might reveal molecular processes underlying the observed length-driven transcriptome imbalance. Within our own experimental data of male C57BL/6 mice housed under specific pathogen free conditions (Fig. 1A,B), those genes (Table S7) enrich for roles in RNA binding, transcription, and splicing. The 1^st^ and 2^nd^ strongest associations are *Neuroblast differentiation-associated protein AHNAK* (AHNAK) and *fused in sarcoma* (Fus), respectively. The former gene was initially identified to encode for an unusually large 700-kDa protein^21^, which was since shown to compete for its expression with a short 17-kDa protein isoform of AHNAK^22^. Fus is a partner^23^ of the polyfunctional age-associated^23–28^ *splicing factor proline and glutamine rich* (Sfpq), which has the 27^th^ strongest association and was recently found to be essential to the transcriptional elongation of genes than 100 kb^6,29^ (Fig. 5A). Our finding that impaired transcriptional efficiency disproportionally disfavoring long transcripts is thus consistent with prior literature^30–34^ and provides a molecular association to the length-driven transcriptional imbalance.

**Fig. 5.**
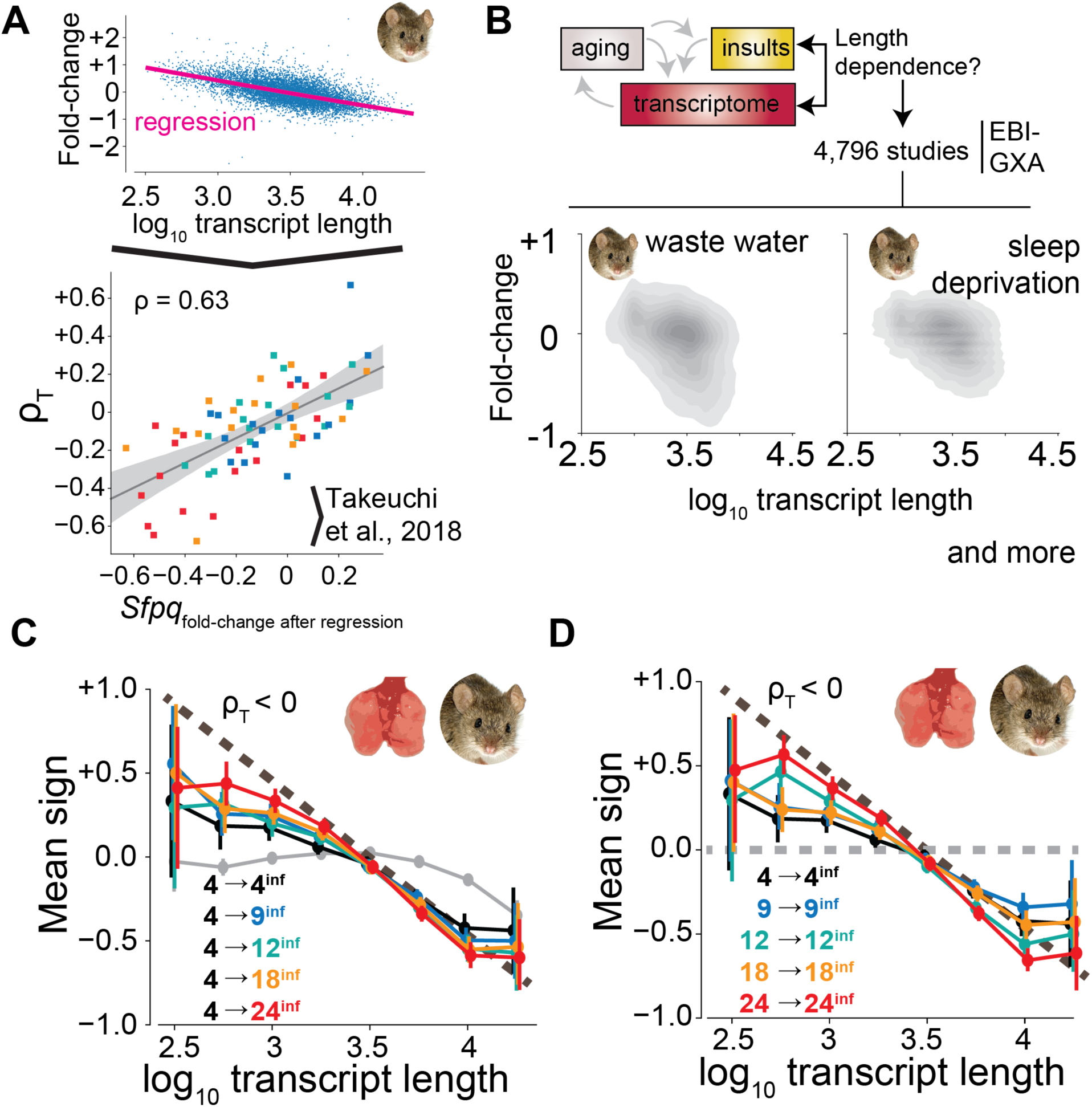
Insults promote length-driven transcriptome imbalance. (**A**) The dependence of gene expression fold-change on gene length, here shown for kidneys from 4-versus 24-month-old mice, can be used to correct the fold-change of specific genes. In the bottom panel, we plot corrected fold-changes of *splicing factor proline and glutamine rich* (*Sfpq*) versus overall transcriptome imbalance as measured by ρ_T_. Squares represent individual organs. Colors represent age as in Fig. 1C. Light grey area shows 95% confidence interval of bootstrapped linear fit. (**B**) Survey of the EBI–Gene Expression Atlas (GXA) for environmental conditions for inducers of transcriptome imbalance. Shown is direction of fold-change for transcripts following exposure to influenza A virus compared with (**C**) 4-month-old uninfected mice or (**D**) age-matched uninfected mice. An average sign of +1 would indicate that all genes are upregulated, whereas an average sign of −1 would indicate that all are downregulated. Dashed brown line is linear approximation of age-dependent trend identified in Fig. 4A. In all cases, we find that influenza promotes a length-driven transcriptome imbalance with a negative slope (ρ < 0). Grey line in (**C**) represents transcriptomes of uninfected lungs.

As environmental factors contribute to aging^35^, we surmise that environmental insults may promote length-driven transcriptome imbalance. Consequently, we perform a meta-analysis of 2,155 mouse and 2,641 human transcriptomic studies represented in the EBI-GXA database, which includes gene expression data for multiple species under different biologic conditions (Fig. 5B). This approach recovers an anticorrelation between transcript length and fold-changes inflicted by inhibitors of transcriptional elongation^31,36^ or exposure to several environmental factors contributing to phenotypic manifestations of aging, such as exposure to pollution^37^, sleep deprivation^38^, heat^39^, and pathogens^14^. Furthermore, we observe an anticorrelation among neurodegenerative disorders such as amyotrophic lateral sclerosis (ALS) and Alzheimer’s disease (Fig. 5B, Tables S8 and S9).

To determine whether those environmental factors could suffice to trigger transcriptome imbalance, we infected mice with a clinically important pathogen in aging^40^, influenza A virus, since its primary target, the lung, is one the two organs that did not yield detectable transcriptome imbalance in 24-month-old mice and elderly humans at the organ level (Fig. S19). In strong support of our hypothesis, we find that influenza A infection causes a downregulation of long transcripts and an upregulation of short transcripts throughout the lifespan of mice, with the effects being strongest in 24-month-old mice (Fig. 5C,D and Fig. S20), the oldest age tested. We conclude that environmental insults that have already been associated with aging^40^ can trigger transcriptome imbalance, and that old individuals may have a reduced capacity to counter environmentally inflicted length-driven transcriptome imbalance.

Length-driven transcriptome imbalance differs from individual gene-specific regulatory mechanisms by being more informative on transcriptome-wide changes observed during aging. It further differs by inherently and preferentially modulating distinctive processes important to the biology of aging. Finally, transcriptome imbalance differs by causing comparatively subtle changes to the transcript levels of individual genes. Conceptually, this matches the notion that manifestations of aging can be observed among living individuals, whereas strong perturbations of essential processes can cause immediate lethality. Moreover, transcriptome imbalance may help to explain the compounding multimorbidity encountered in elderly humans^41^, and why qualitatively distinct insults funnel into similar phenotypes during aging^1^. Our study further opens fundamental questions on the organization of transcriptomes and the interplay between epigenetic, transcriptional, and proteomic homeostasis^42,43^.

## Acknowledgments

We thank Ermelinda Ceco, Ching-I Chen, Cong Chen, Yuan Cheng, Monica Chi, Stephen Chiu, Carla Cuda, Annette Flozak, Francisco Gonzales, Katie Helmin, Erin Hogan, Philip Homan, Dana Kimelman, Emilia Lecuona, Natalia Mangani, Vince Morgan, Trevor Nicholson, Rojo Ratsimandresy, Constance Runyan, Rana Saber, and Saul Soberanes for sample preparation; Elizabeth Bartom for the template of bioinformatic processing of the RNA sequences toward counts; Paul A. Reyman and Rohan Verma for creating the template for differential gene expression; Chao Wang and Salvatore Loguerico for comments on the manuscript; Jennifer Davis for proofreading; and Northwestern University IT and Genomics Nodes for infrastructural support.

## Author contributions

T.S. conceptualized the data analysis, discovered length-driven transcriptome imbalance, performed data curation, developed software, performed the data analysis, visualized results, cowrote the original draft, and supervised the upscaling of bioinformatic preprocessing; R.A.G. performed pilot differential expression queries in GTEx; A.C.M. performed experiments; K.A., S.S.L., and H.T.N. upscaled bioinformatic preprocessing; B.S. performed experiments; H.A.V. performed RNA sequencing; M.S. and M.P.T. performed experiments; H.P. supervised parts of the mouse experiments; W.E.B., N.C., and K.R. supervised parts of the experiments and reviewed the manuscript; J.I.S. reviewed and edited the manuscript and acquired funding; R.I.M. supervised the study, reviewed and edited the manuscript, and acquired funding; A.V.M. conceptualized, supervised, and performed mouse experiments and supervised development of bioinformatic templates; G.R.S.B. conceptualized the study, administrated the project, acquired funding, and reviewed and edited the manuscript; and L.A.N. conceptualized and supervised data analysis and cowrote the original draft.

## Data and materials availability

RNA sequencing data created during this study will be shared on the Sequence Read Archive upon publication.

## Materials and Methods

### Animals

All mouse procedures were approved by the Institutional Animal Care and Use Committee at Northwestern University (Chicago, IL, USA). All strains including wild-type mice are bred and housed at a barrier and specific pathogen–free facility at the Center for Comparative Medicine at Northwestern University. Male C57BL/6 mice were provided by NIA NIH and were housed at Northwestern University Feinberg School of Medicine Center for Comparative Medicine for 4 weeks prior to sacrifice.

Mice were euthanized by pentobarbital sodium overdose. Immediately the chest cavity was opened and animals were perfused via the left ventricle with 10 mL of HBSS (Ca/Mn+). The following organs were harvested: lung, heart, liver, kidney, adrenal gland, white (perigonadal) and brown adipose tissue, skin, skeletal muscles, frontal cortex, cerebellum, esophagus, stomach, and small and large intestine. Gut epithelial cells were isolated after flushing large intestine with EDTA/EGTA solution. Lung and skeletal muscle were subjected to enzymatic digestion to release stromal and immune cells and sorted by fluorescence-activated cell sorting as described elsewhere ^44^. All tissues and cells were immediately frozen on dry ice and stored at −80°C for processing.

### RNA isolation and RNA sequencing

RNA was isolated using an RNeasy DNA/RNA kit after homogenization and lysis in guanidine thiocyanate buffer supplemented with β-mercaptoethanol. RNA concentration and quality were assessed using an Agilent TapeStation. RNAseq libraries were prepared using an NEB Next RNA Ultra kit with polyA enrichment module using an Agilent Bravo NGS Automated fluidics handling platform as described elsewhere^44^. Libraries were multiplexed and sequenced on an Illumina NextSeq 500 platform using 75 cycles of high-output flow cells and a dual indexing strategy.

### Bioinformatics

Sequencing reads were analyzed using an implementation of Ceto (https://github.com/ebartom/NGSbartom) in Nextflow^45^. Briefly, BCL files were demultiplexed and converted to fastq files using bcl2fastq, version 2.17.1.14, with default parameters. The raw reads were trimmed using trimmomatic^46^, version 0.36, with the following parameters: trailing = 30 and minlen = 20. Trimmed reads were aligned to the mouse reference genome (GRCm38.p3) with annotations from Ensembl v78 using tophat, version 2.1.0^47^, with the following parameters: no novel junctions, read-mismatches = 2, read-edit-distance = 2, and max-multihits = 5. Aligned reads were counted using Htseq-count from htseq^48^, with the following parameters: intersection-nonempty, reverse strand, feature-type = exons, and id-attribute = gene_id.

For GTEx^13^, count matrices (version 7) were downloaded from GTExPortal. Samples corresponding to cell lines were dismissed from any further analysis.

### Differential expression

Differential gene expression analysis was performed with DESeq2, version 1.20 (mouse) and 1.22^49^ using an α value of 0.05 for the adjusted *P* value cutoff. We subsequently only kept genes that mapped unambiguously been Ensembl Gene Identifiers and NCBI (Entrez) gene identifiers^4^.

### Characteristics of genes

For transcription factors, we mapped the Gene Transcription Regulatory Database v18_06^50^ to ±200 nucleotides to transcriptional start sites supplied by BioMart for the human reference genome build GRCh38.p12 and the mouse reference genome build GrCm38.p6. For miRNAs we used miRDB_v5.0^51^. For mature transcripts, length parameters and CG content were identified from GenBank and mapped to genes as described elsewhere using the median across different transcripts^4^. Number of exons, and their minimal, median, and maximal length, were extracted from BioMart. For genes and chromosomes, characteristics were extracted as described elsewhere^4^.

### Modeling

Gradient boosting regression models were built in scikit-learn, version 0.20.3^3^. Of the transcripts, 90% were included and 10% were used as a test set. We only considered protein-coding genes with at least one research publication and an official gene symbol, and which unambiguously mapped in a 1:1 relation between NCBI (Entrez) gene identifiers and Ensembl Gene Identifiers.

### Kernel-density visualizations

Kernel density visualizations were created with Seaborn (https://github.com/mwaskom/seaborn) using default parameters.

### Symbols

Pictures of killifish, mice, and rats were obtained from wiki-commons, and pictures of humans from the Noun Project and OliM, under creative commons license.

### Reanalysis of prior studies

We considered genes reported to be significantly up- or downregulated in earlier studies. For mice and rats we used protein-coding genes with at least one research publication and an official gene symbol, and the median transcript lengths derived from GenBank. For killifish we used genes and gene lengths as reported by Reichwald et al. 2015^10^.

### Transcriptome imbalance

We defined the extent of the transcriptome imbalance as the Spearman correlation between transcript length and fold-change of transcripts in older individuals over younger ones. Significance was obtained through the scipy.stats, version 1.2.1, implementation of the Spearman correlation^52^.

### Single-cell length correlation

Data of Kimmel et al. ^11^ and Tabula Muris Senis^12^ were downloaded from http://mca.research.calicolabs.com/ and https://figshare.com/articles/Processed_files_to_use_with_scanpy_/8273102, respectively. As cell types we considered the cell_type and cell_ontology_class columns within the respective meta-date tables contained in the h5ad files. For consistency between the two studies, we further renamed “classical monocyte” to “monocyte”, “natural killer cell” to “NK cell”, “Lung endothelial cell” to “endothelial cell”, and “alveolar macrophage” to “macrophage”. We only considered protein-coding genes which were detected in at least one cell of a given cell type in an individual organ in a given study. We determined the single-cell length correlation by measuring the Spearman correlation between transcript length and signal reported by the studies of Kimmel et. al and Tabula Muris Senis, respectively. For Tabula Muris Senis we solely considered the subset of the data corresponding to FACS-isolated single cells as they demonstrated the highest sensitivity according to their study^12^.

### Functional enrichments

We considered the genes with the 5% shortest and 5% longest median transcript length. We obtained the categorization of mutants from HAGR^53,54^ and mapped them to human and mouse orthologues through Homologene, version 68 (https://ftp.ncbi.nlm.nih.gov/pub/HomoloGene). We considered genes labeled as pro-longevity to be beneficial, and genes labeled as anti-longevity to be deleterious. For Gene Ontologies we used the mapping to NCBI provided by the National Library of Medicine (https://ftp.ncbi.nlm.nih.gov/gene/DATA/gene2go.gz) and considered any nonnegating annotation. For differential enrichment we considered genes enriched among the genes with transcripts of one length extreme (5% shortest and 5% longest median) at a Benjamini-Hochberg *P* value of <0.05 and depleted among the genes with the other length extreme.

### Annotation network construction

Annotations were represented as nodes, and we drew edges between them if at least one gene carried either annotation and had been identified by the preceding enrichment analysis. Subsequently we trimmed edges. First, we kept those edges where the largest fraction of the genes of the smaller node were included in the larger. Second, for a given pair of the smaller and larger node, we kept the link if the larger node was the smallest larger node connected to the smaller node. Third—if there were still multiple distinct edges for a smaller node—we kept those where overall there would be fewer genes annotated for the larger node (irrespective of number observed in enrichment analysis).

### Identification of genes correlating with transcriptome imbalance

First, the global relation between transcript length and fold-change was approximated through a Lowess fit using Statsmodels, version 0.9^55^. Second, we defined residual fold-changes by subtracting the Lowess fits from the observed fold-changes. For mice, we considered the differential gene expression analyses of our own census of mice between 4-month-old and 9-, 12-, 18-, or 24-month-old mice. For humans, we considered GTEx differential gene expression analyses between donors in their 2nd decade and donors in their 4th, 5th, 6th, or 7th decade, as well as between donors in their 3rd decade and donors in their 4th, 5th, 6th, or 7th decade (hence yielding up to eight comparisons per gender and subregion). We defined the correlation between gene and transcriptome imbalance as the Spearman correlation between transcriptome imbalance and the residual fold-changes.

### Comparison to EBI-GXA

We downloaded the EBI-GXA^7^ in spring 2017^4^. We parsed the experimental descriptions from the config files supplied and the fold-changes from the supplied differential expression reports. We only considered protein-coding genes with at least one research publication and an official gene symbol, and which unambiguously mapped in a 1:1 relation between NCBI (Entrez) gene identifiers and Ensembl Gene Identifiers.

### Influenza A virus infection

Influenza virus strain A/WSN/1933 (WSN) was grown for 48 h at 37.5°C and 50% humidity in the allantoic cavities of 10–11-d-old fertile chicken eggs (Sunnyside Hatchery, WI). Viral titers were measured by plaque assay in Madin-Darby canine kidney epithelial cells (American Type Culture Collection). Virus aliquots were stored in liquid nitrogen and freeze/thaw cycles were avoided. For infection, mice were anesthetized with isoflurane and infected intratracheally with 150 plaque forming units (PFU) in 50 μL of PBS. Mice were sacrificed after 4 days.

**Fig. S1.**
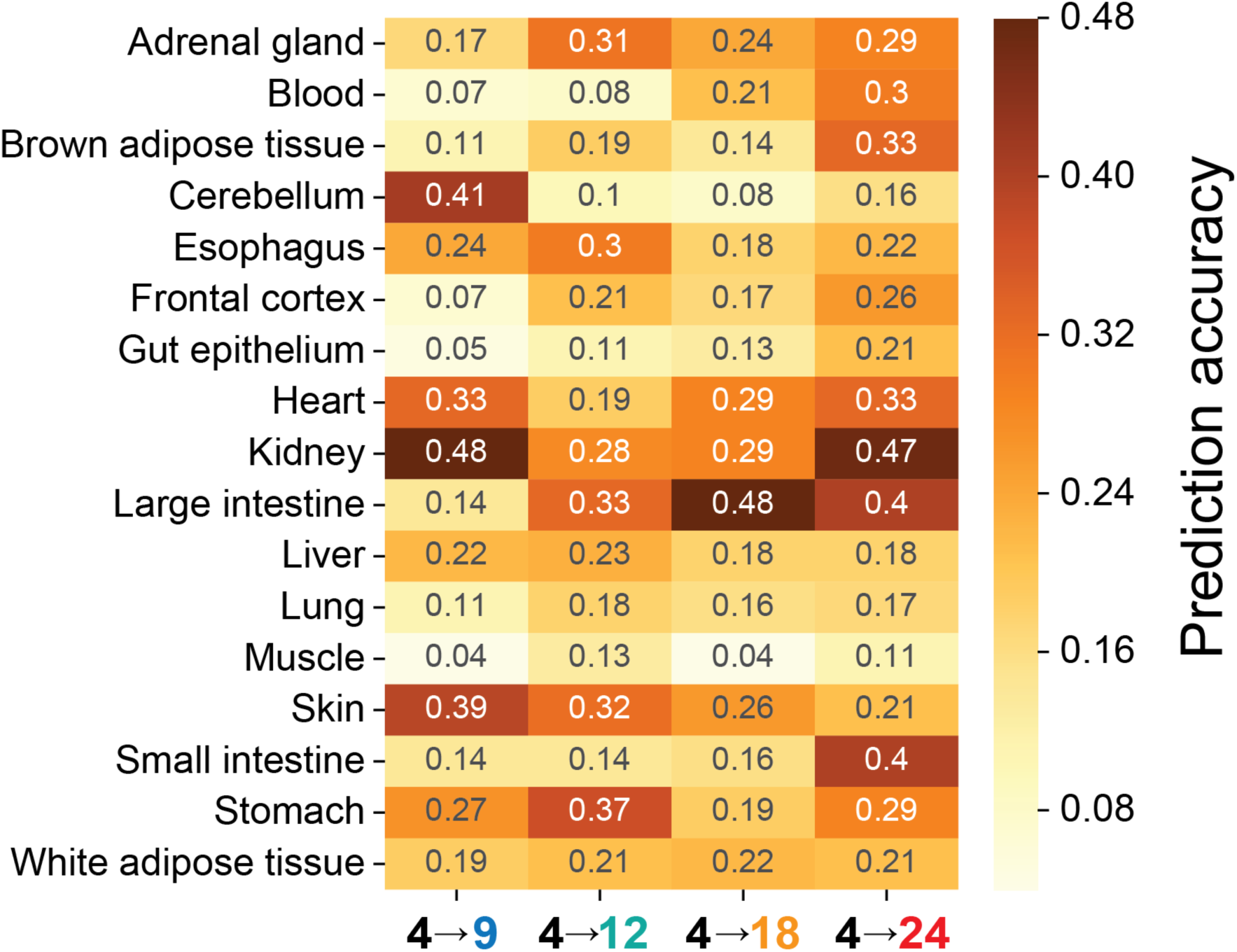
Prediction accuracy. Prediction accuracy was defined as the Spearman correlation between observed and predicted fold-changes.

**Fig. S2.**
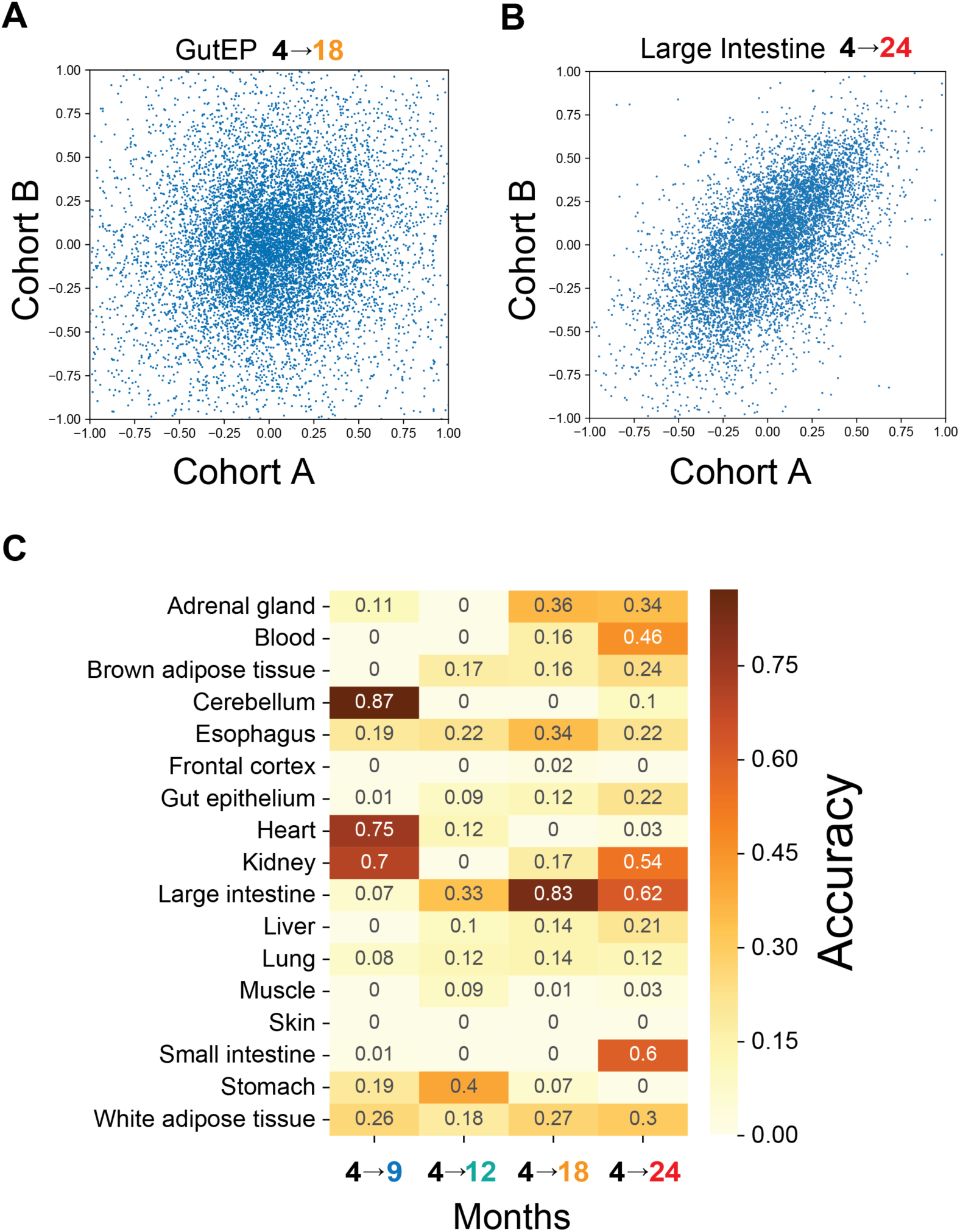
Replicability of fold-changes among cohorts. (**A**) Fold-changes of 18-month-old gut epithelium over 4-month-old gut epithelium. (**B**) Fold-changes of 24-month-old large intestine over 4-month-old large intestine. (**C**) Empirically observed accuracy was defined as the Spearman correlation between the fold-changes of both cohorts.

**Fig. S3.**
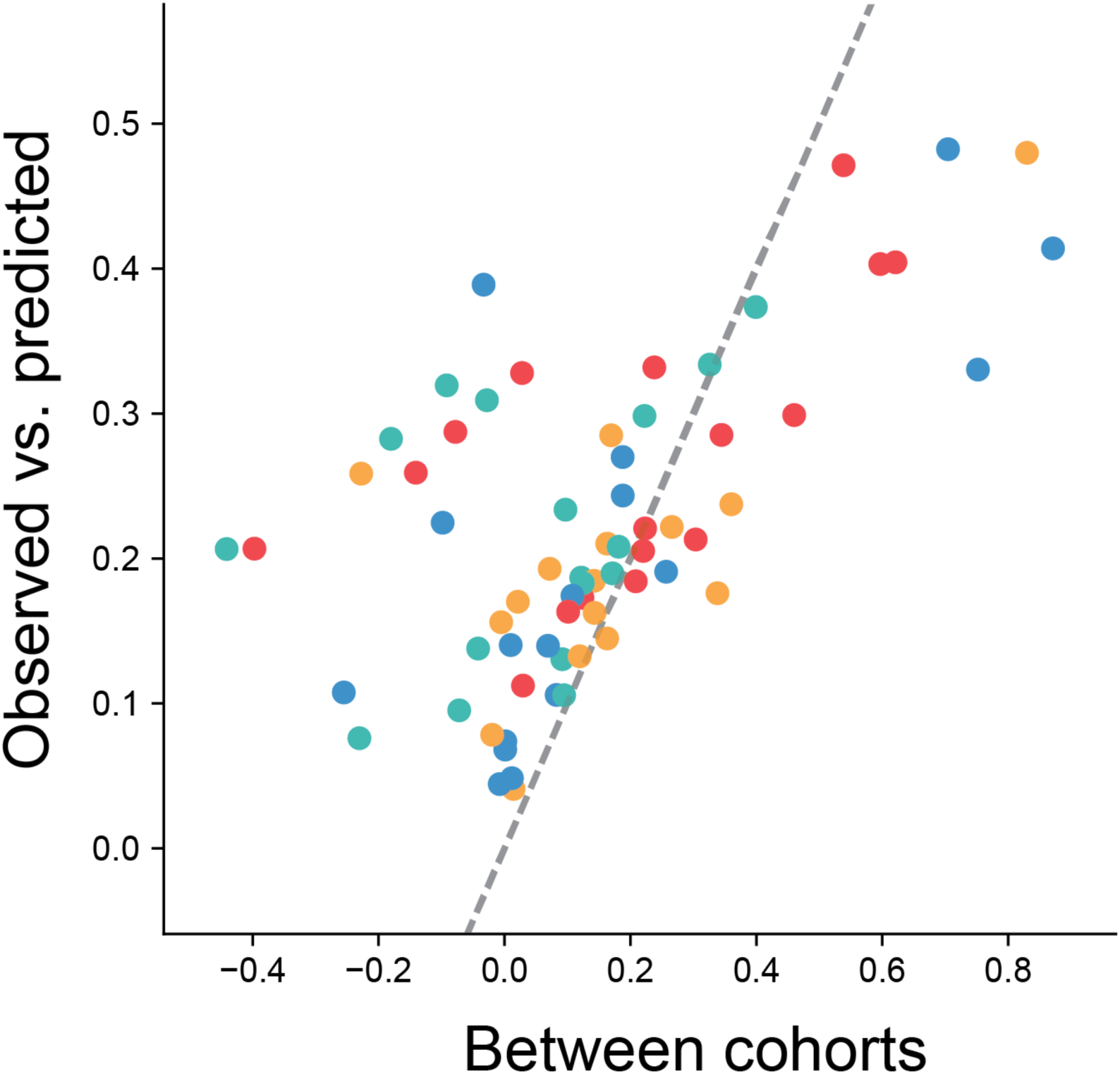
Comparison of prediction accuracy and empirically observed accuracy. Comparison of empirically observed accuracy between cohorts (Fig. S2) and accuracy of predictions (Fig. S1). Colors indicate ages as in Fig. 1A.

**Fig. S4.**
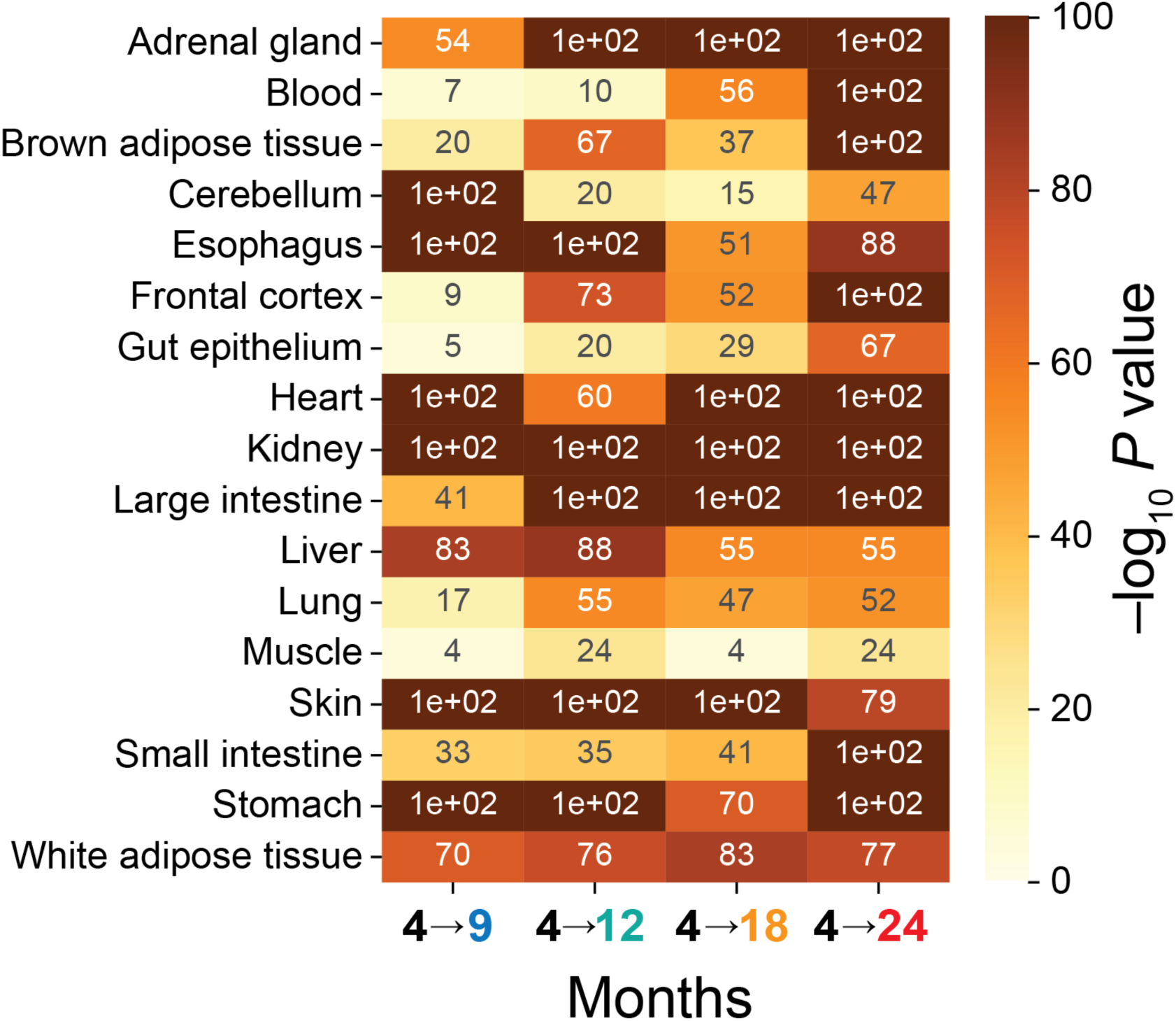
Significance of prediction. Significance of Spearman correlation between observed and predicted fold-changes reported in Fig. S1.

**Fig. S5.**
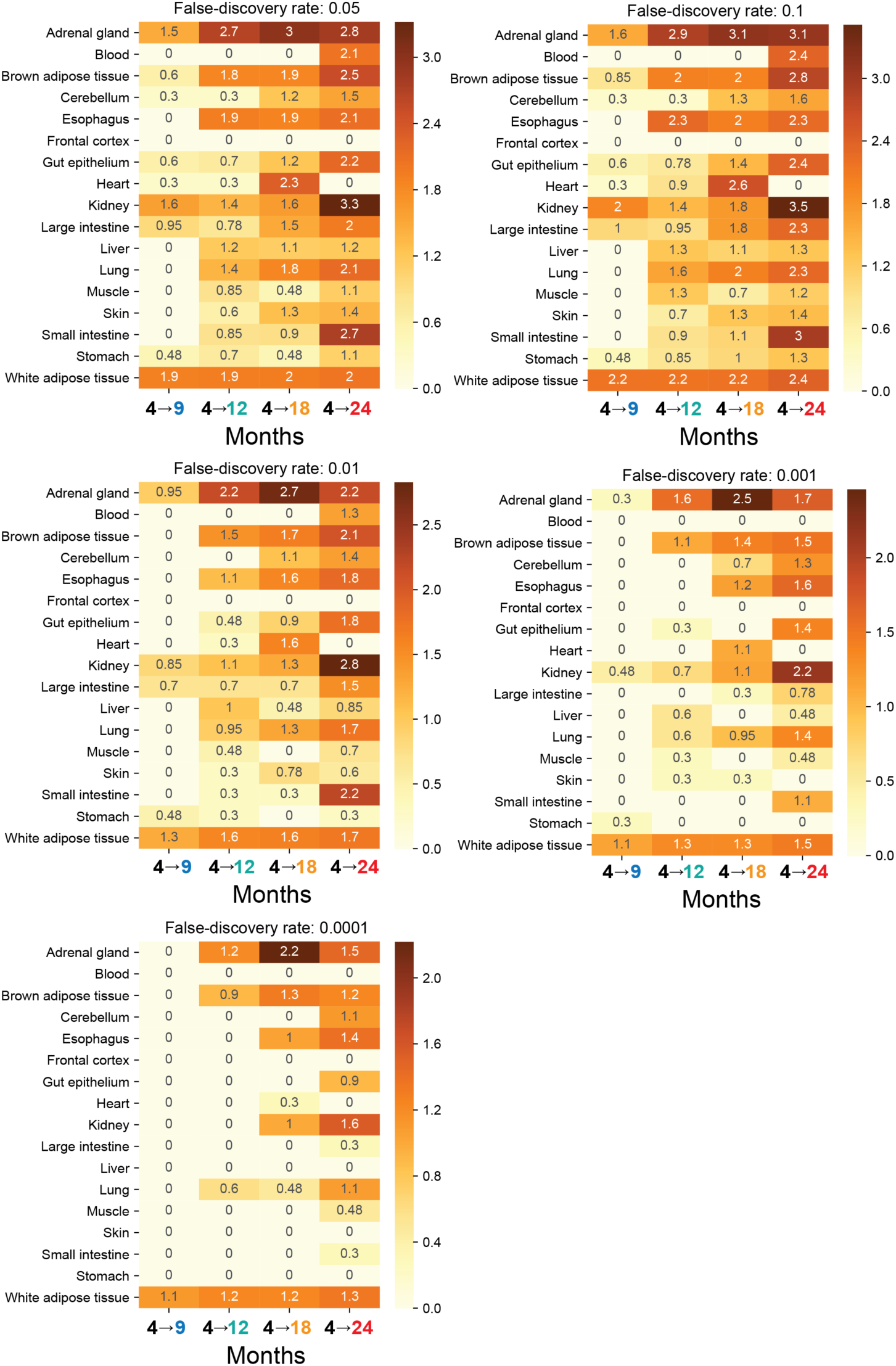
Differentially expressed genes. Number of differentially expressed genes relative to 4-month-old organs at different thresholds for false discovery rate (0.1, 0.05, 0.01, 0.001, and 0.0001). Values are log_10_(differentially expressed genes + 1).

**Fig. S6.**
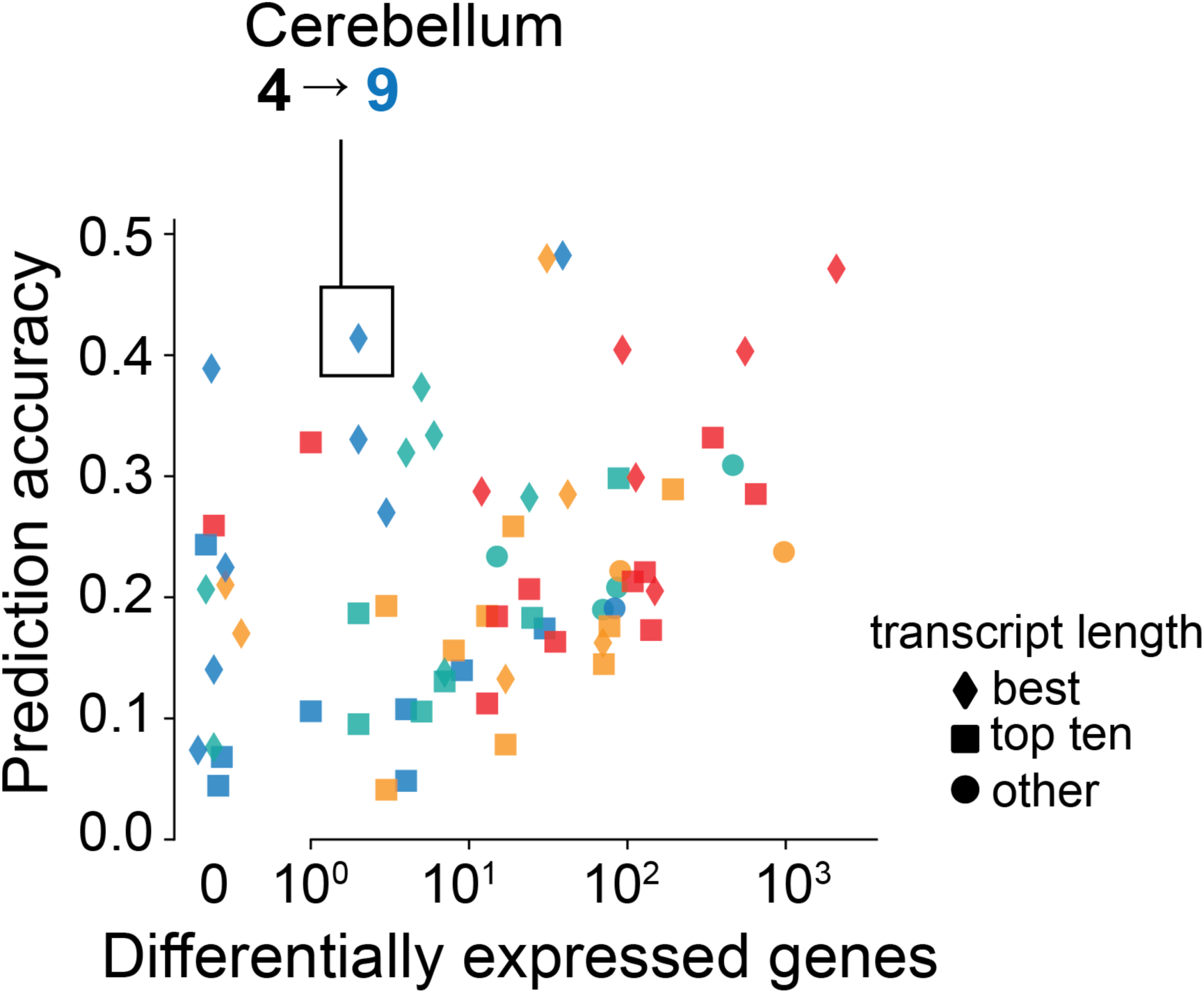
Comparison between prediction accuracy and differential expression. Prediction accuracy, defined as Spearman correlation between observed and predicted fold-change for individual organs of our study, is compared with the number of genes differentially expressed at a false discovery rate of 0.05. Colors represent age as in Fig. 1A, and symbols indicate ranking of transcript length (one of multiple length-related features) within age- and organ-specific models.

**Fig. S7.**
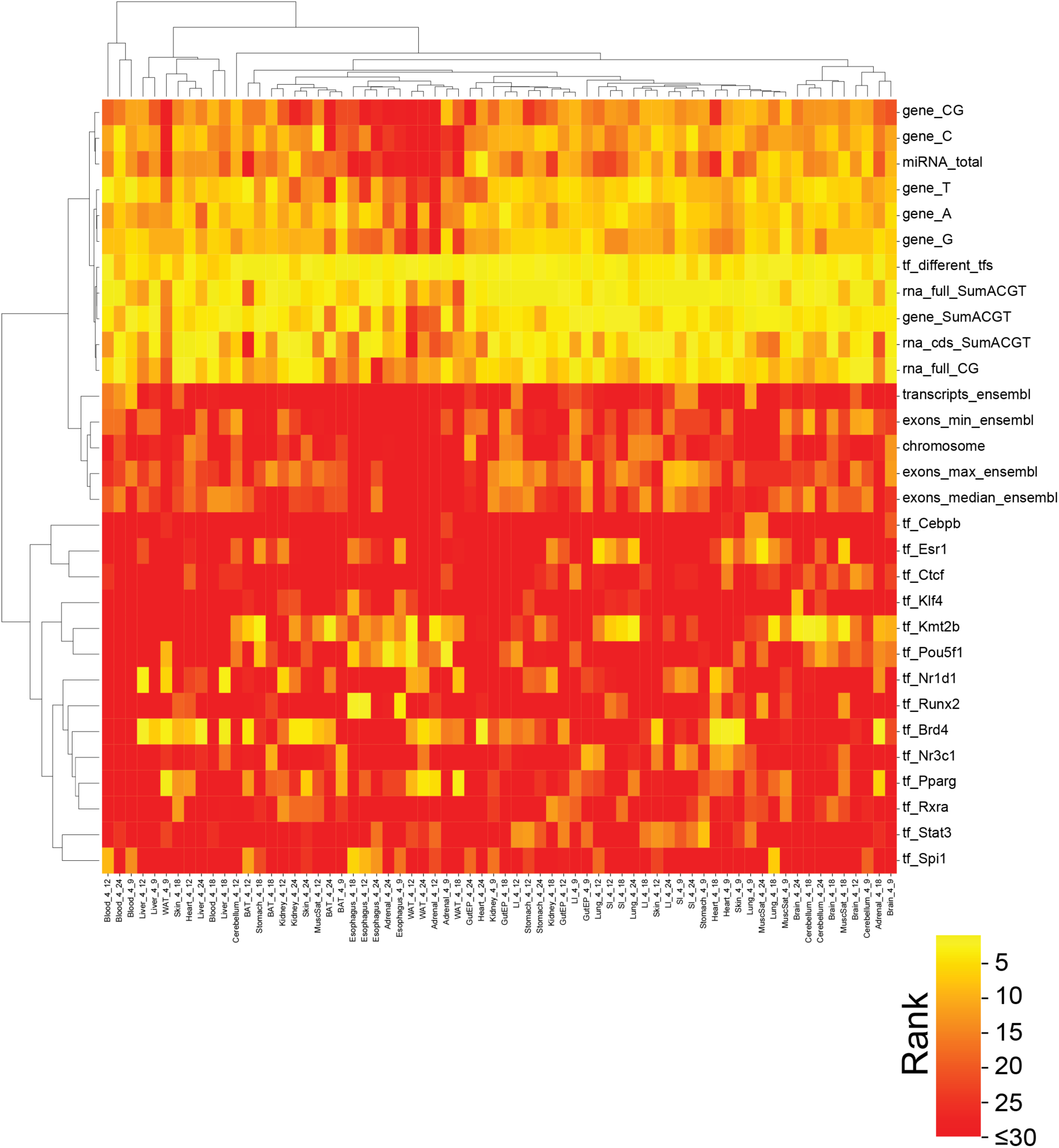
Cluster map of most informative features in mice. Most informative features (median rank across organs and ages lie in top 30) grouped by Ward clustering.

**Fig. S8.**
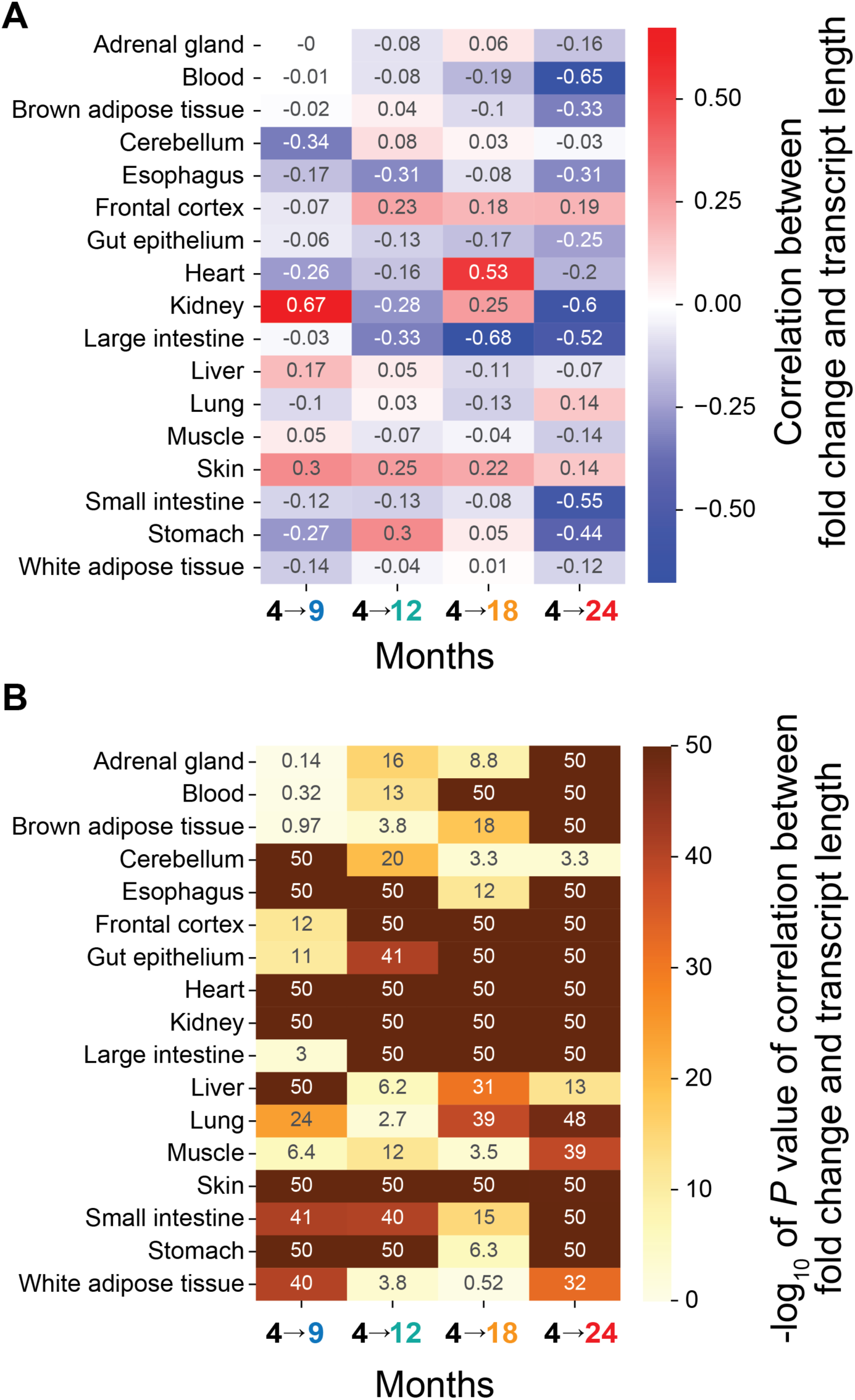
Transcript length and fold-changes. (**A**) Spearman correlation between median transcript length and observed fold-changes. (**B**) Significance of correlations.

**Fig. S9.**
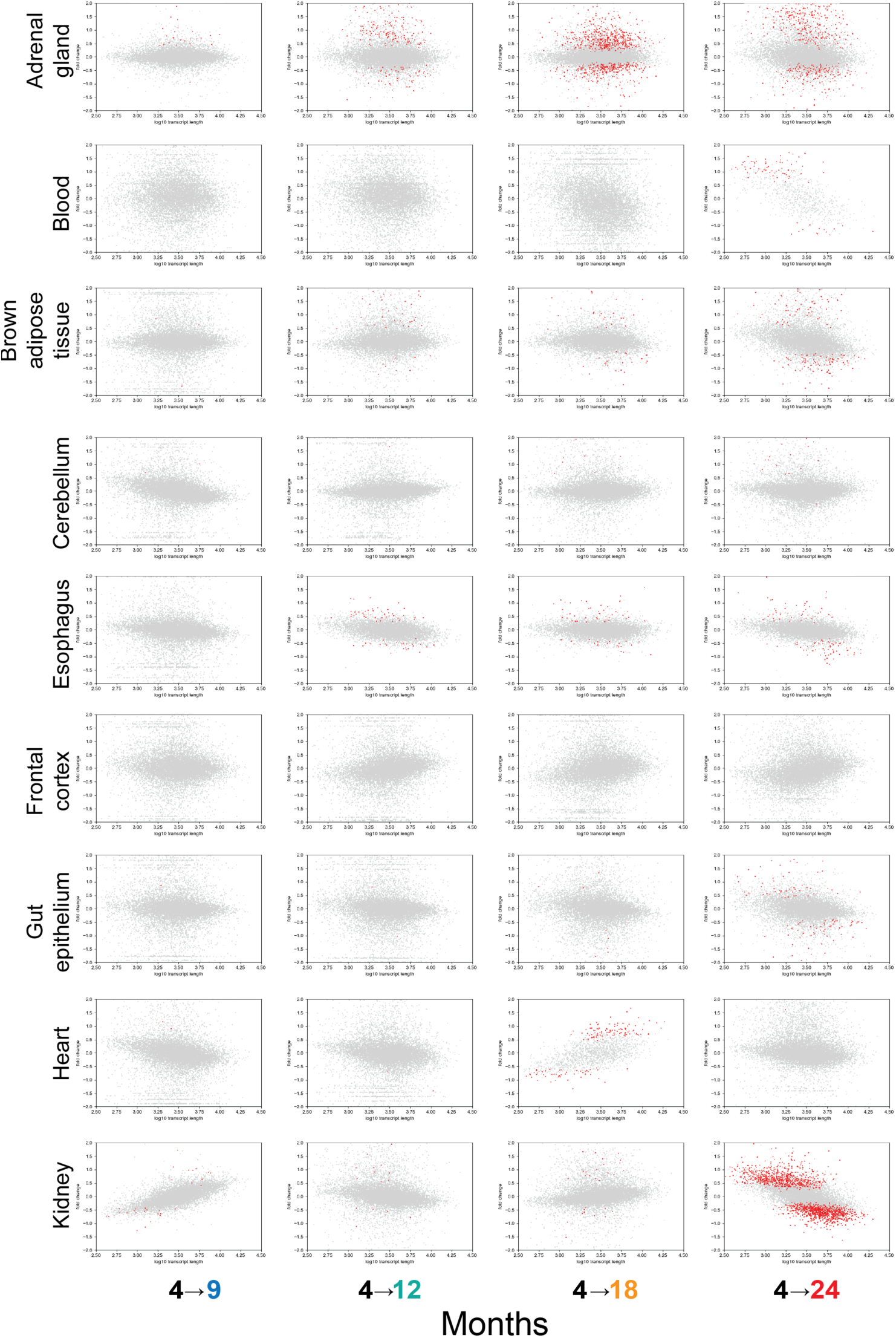
Organ-specific representation of transcript length and fold-changes, part 1. Comparison for adrenal gland, blood, brown adipose tissue, cerebellum, esophagus, frontal cortex, gut epithelium, heart, and kidney. Grey dots are genes. Red dots are genes identified to be differentially expressed at a false discovery rate of 0.05.

**Fig. S10.**
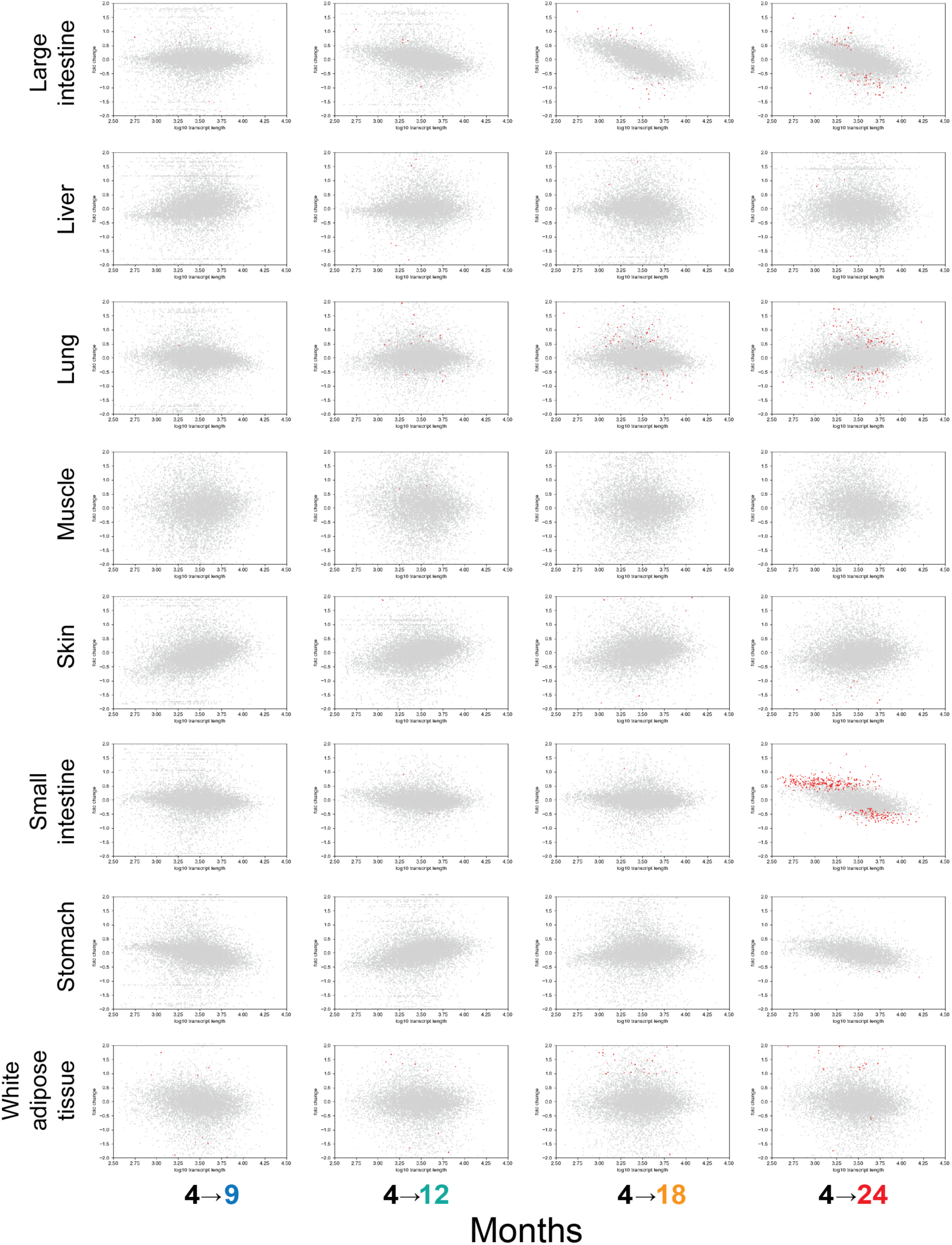
Organ-specific representation of transcript length and fold-changes, part 2. Comparison for large intestine, liver, lung, muscle, skin, small intestine, stomach, and white adipose tissue. Grey dots are genes. Red dots are genes identified to be differentially expressed at a false discovery rate of 0.05.

**Fig. S11.**
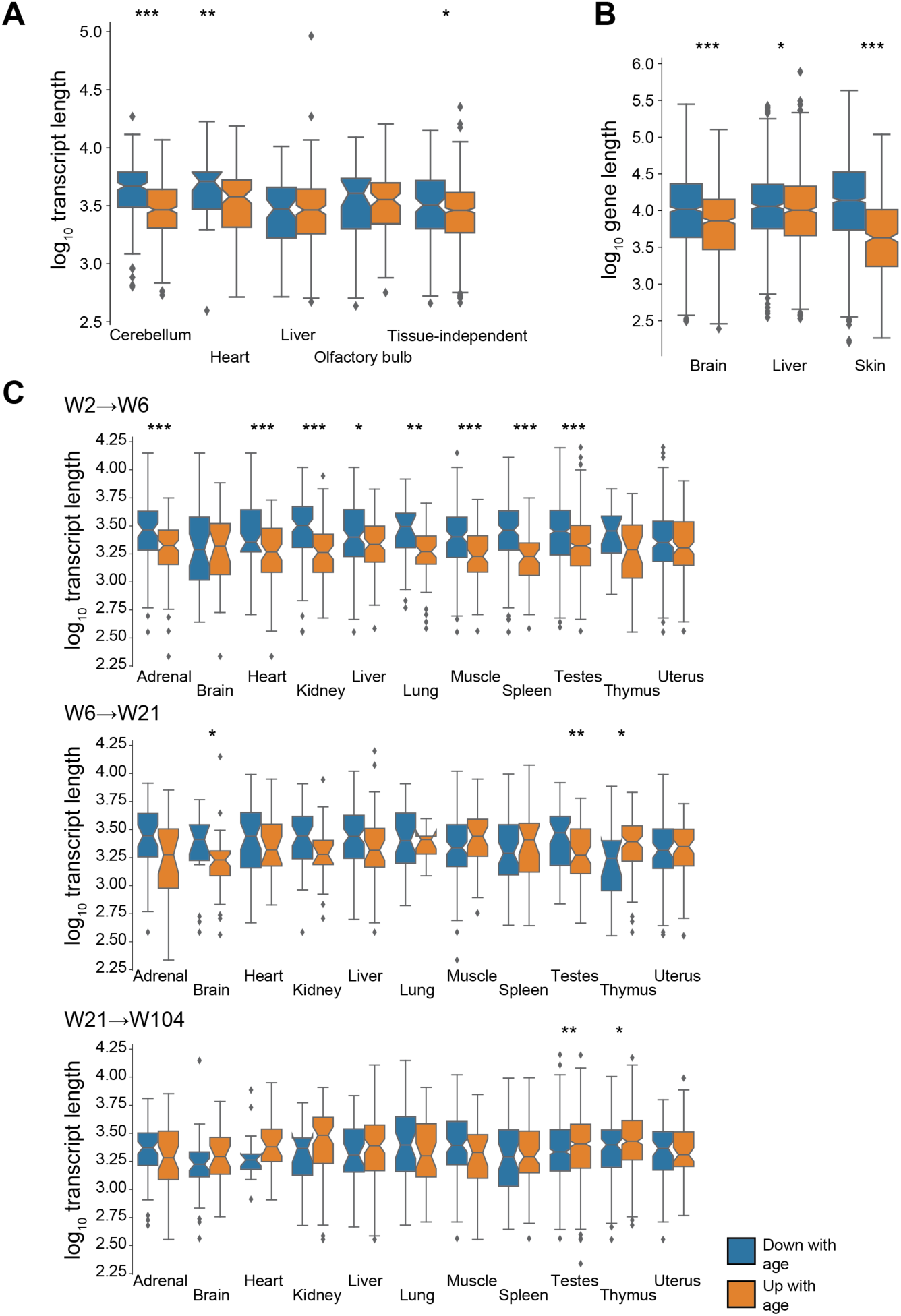
Differential length of down- and upregulated genes. (**A**) Median transcript length for mouse genes reported to be differentially expressed across 3-, 12-, and 29-month-old animals by Benayoun et al. 2019^5^. (**B**) Median gene length for killifish genes reported to be differentially expressed between 5 and 39 weeks of age by Reichwald et al. 2015^10^. Gene lengths are as reported. (**C**) Median transcript length for rat genes reported to be differentially expressed by Yu et al. 2014^9^. W2, W6, W21, and W104 indicate weeks after birth. \**P* < 0.05, \*\**P* < 0.01, and \*\*\**P* < 0.001 in two-sided Mann-Whitney *U* test.

**Fig. S12.**
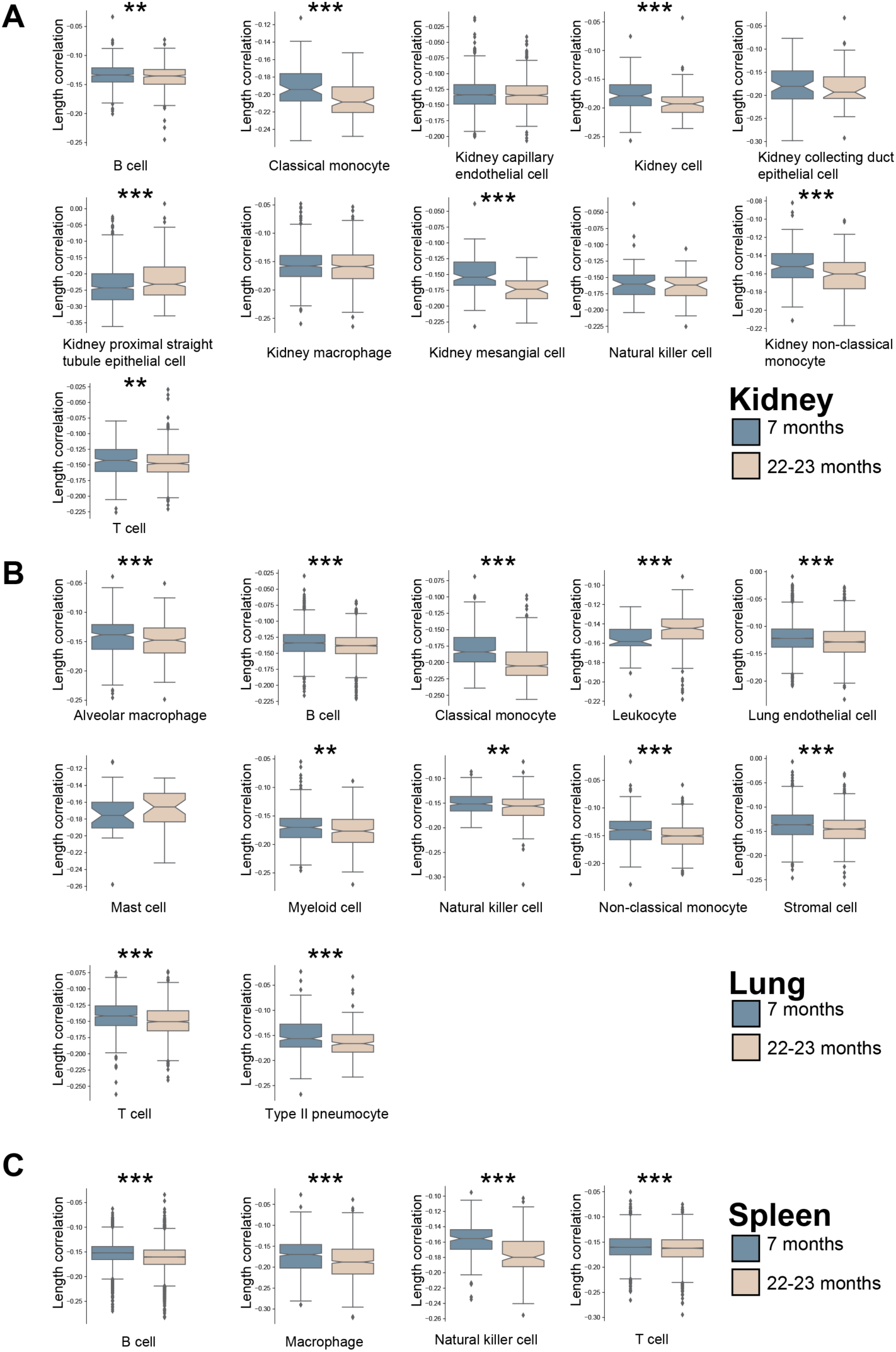
Differential correlation between transcript length and transcript counts in single cells. Single cells of indicated cell types of 7- and 22-23-months old mice^11^ for (**A**) Kidney (**B**) Lung (**C**) Spleen. \**P* < 0.05, \*\**P* < 0.01, and \*\*\**P* < 0.001 in two-sided Mann-Whitney *U* test.

**Fig. S13.**
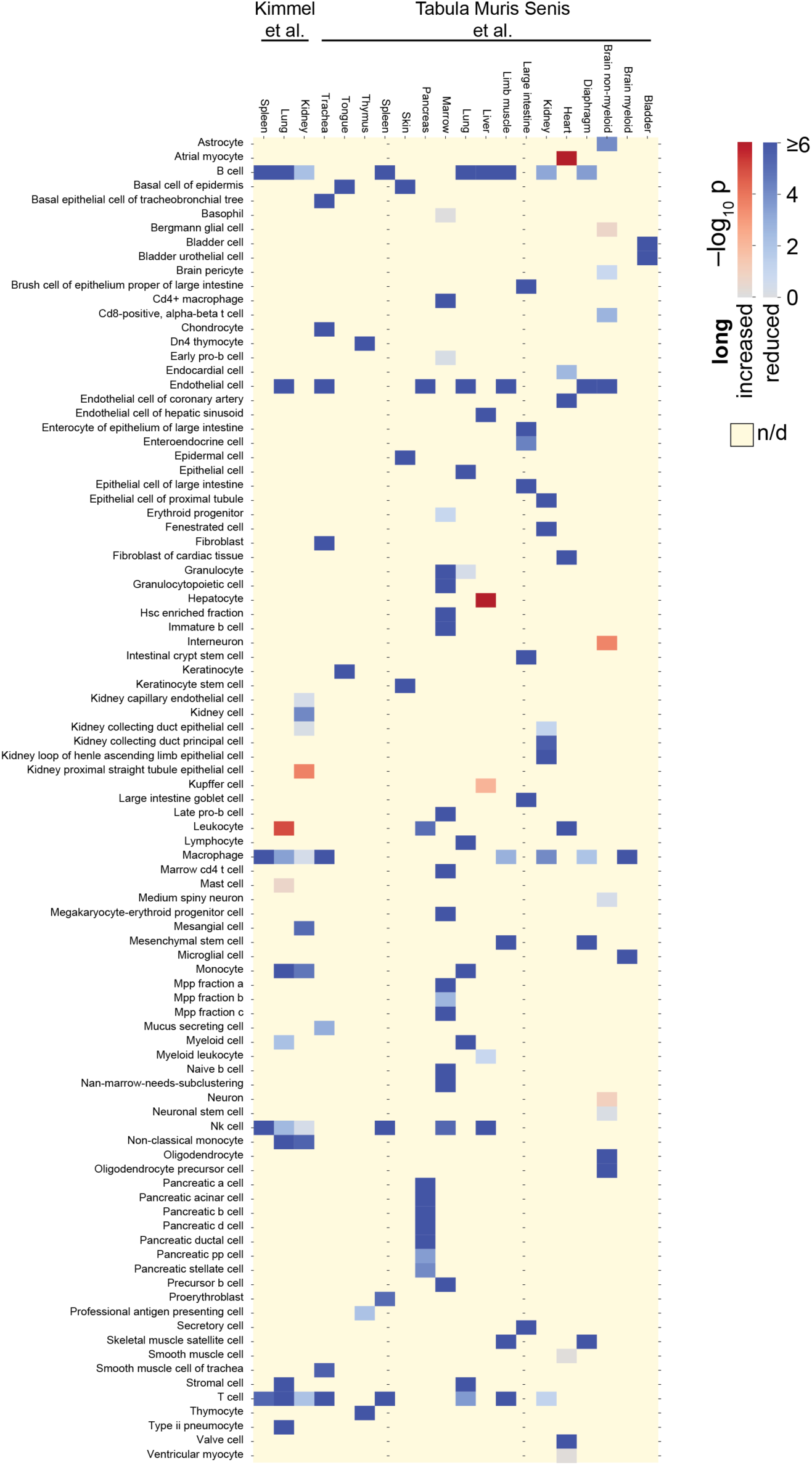
Significance of difference in single-cell length correlations. As in Fig. 2C, but with labels of cell types assigned by authors^11,12^.

**Fig. S14.**
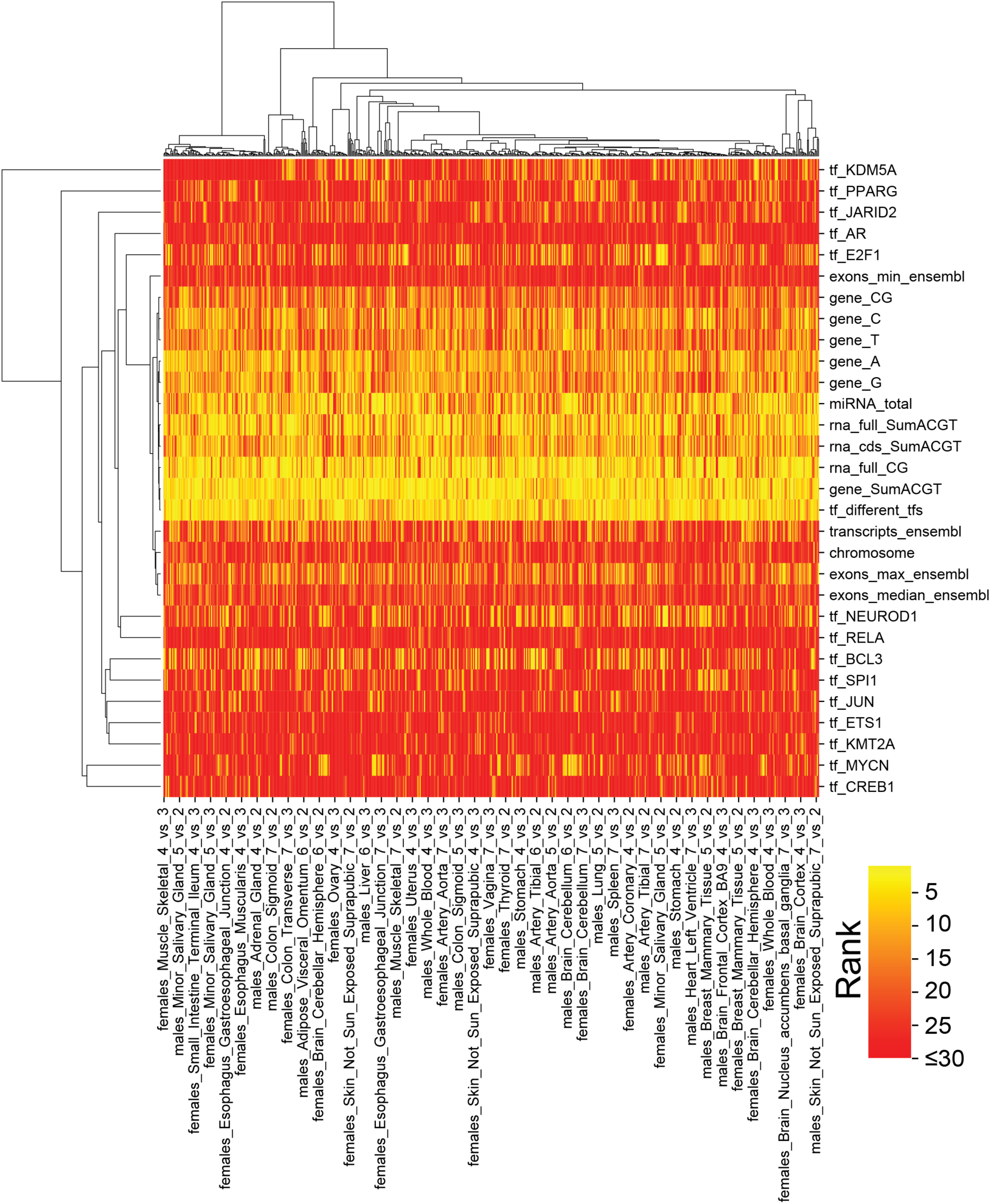
Cluster map of most informative features in human GTEx. Most informative features (median rank across organs and ages lie in top 30) grouped by Ward clustering.

**Fig. S15.**
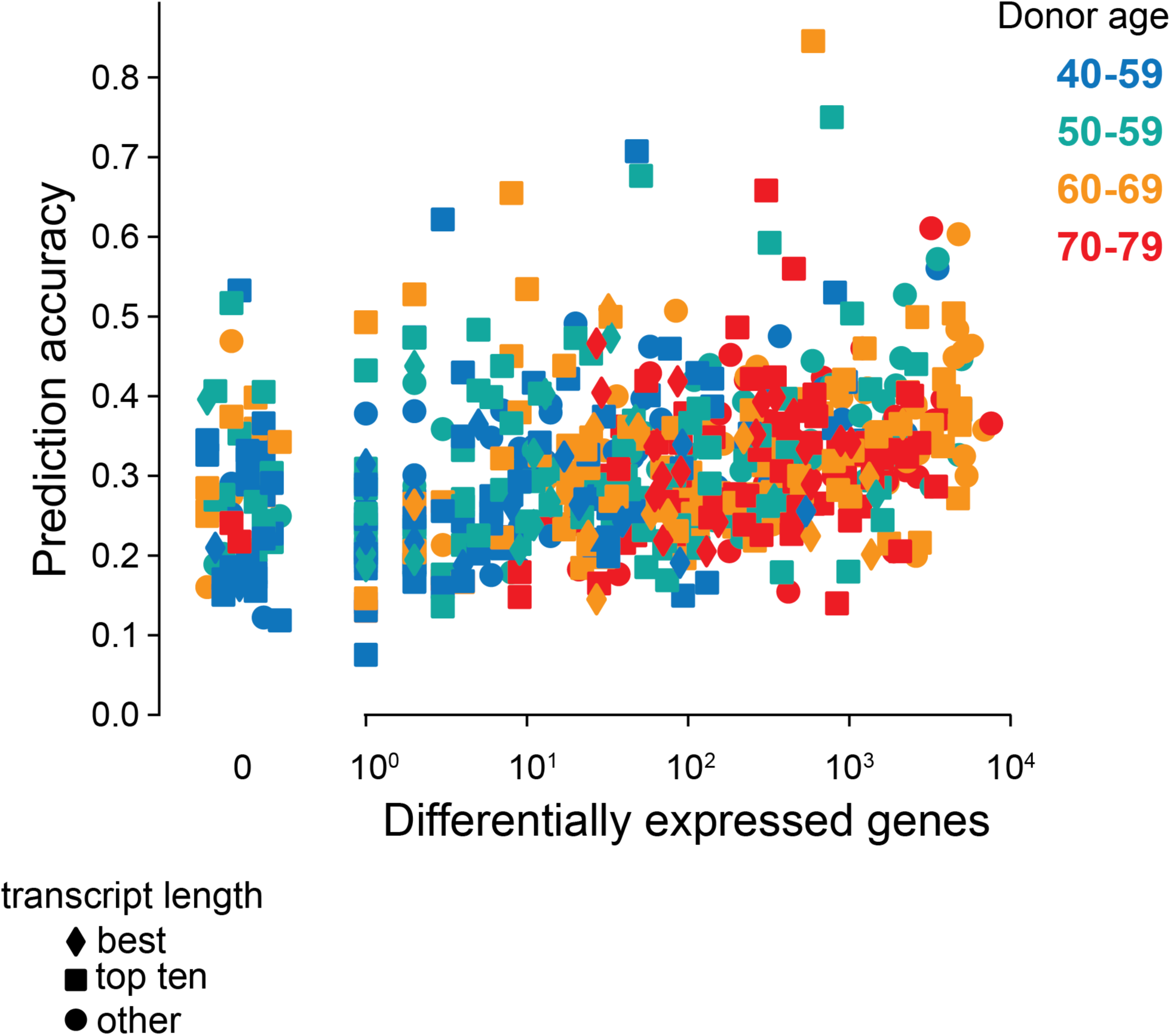
Prediction accuracy and number of differentially expressed genes for human GTEx. Analogous to Fig. S6, but for human GTEx samples. Shown are comparisons between donors in the indicated decade relative to donors aged 20–29 years. Male and female donors are represented separately.

**Fig. S16.**
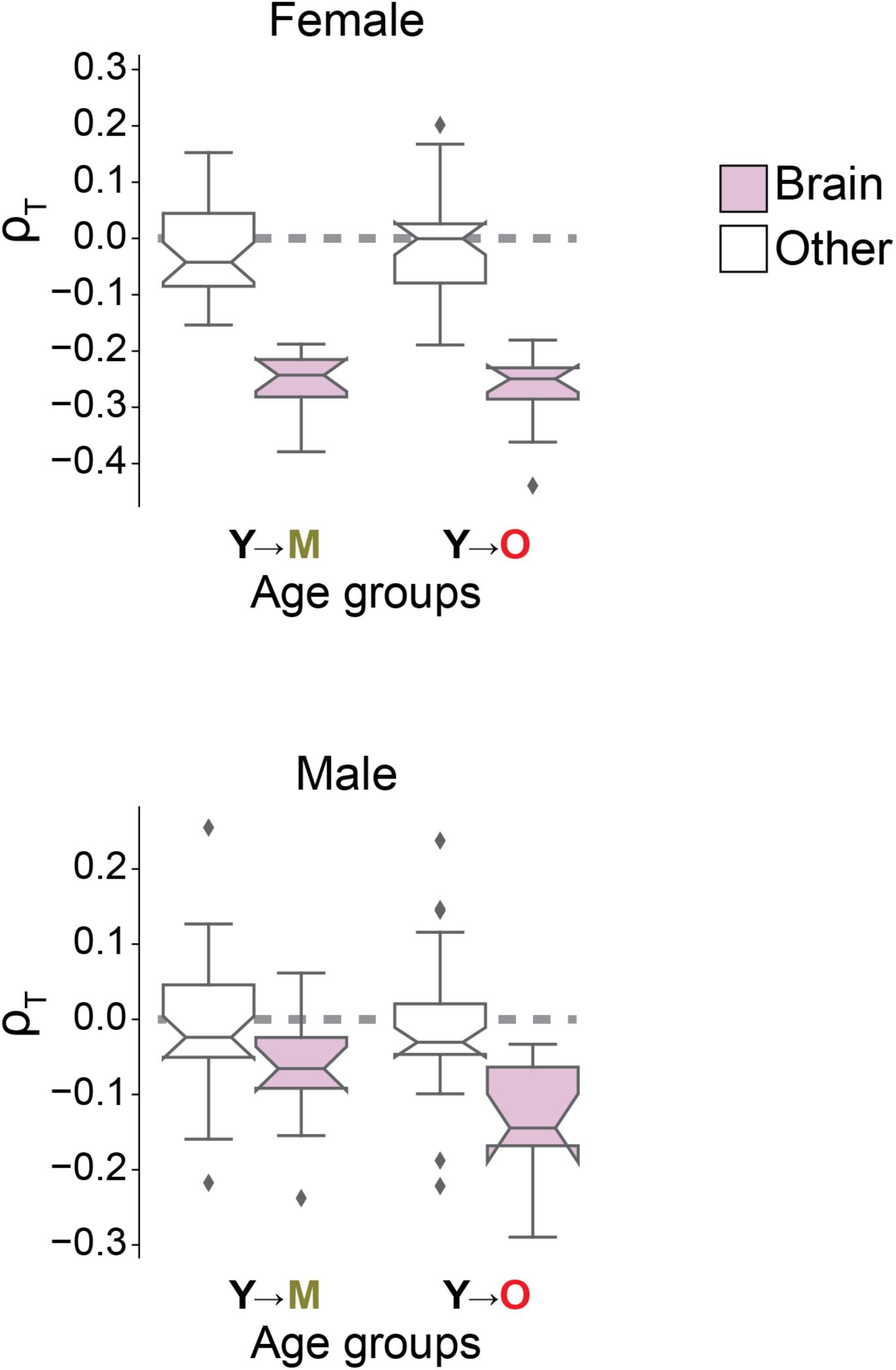
Gender-specific imbalance among human donors. As Fig. 3C, but displaying length-driven transcriptome imbalance separately for tissues of female (top) and male donors (bottom).

**Fig. S17.**
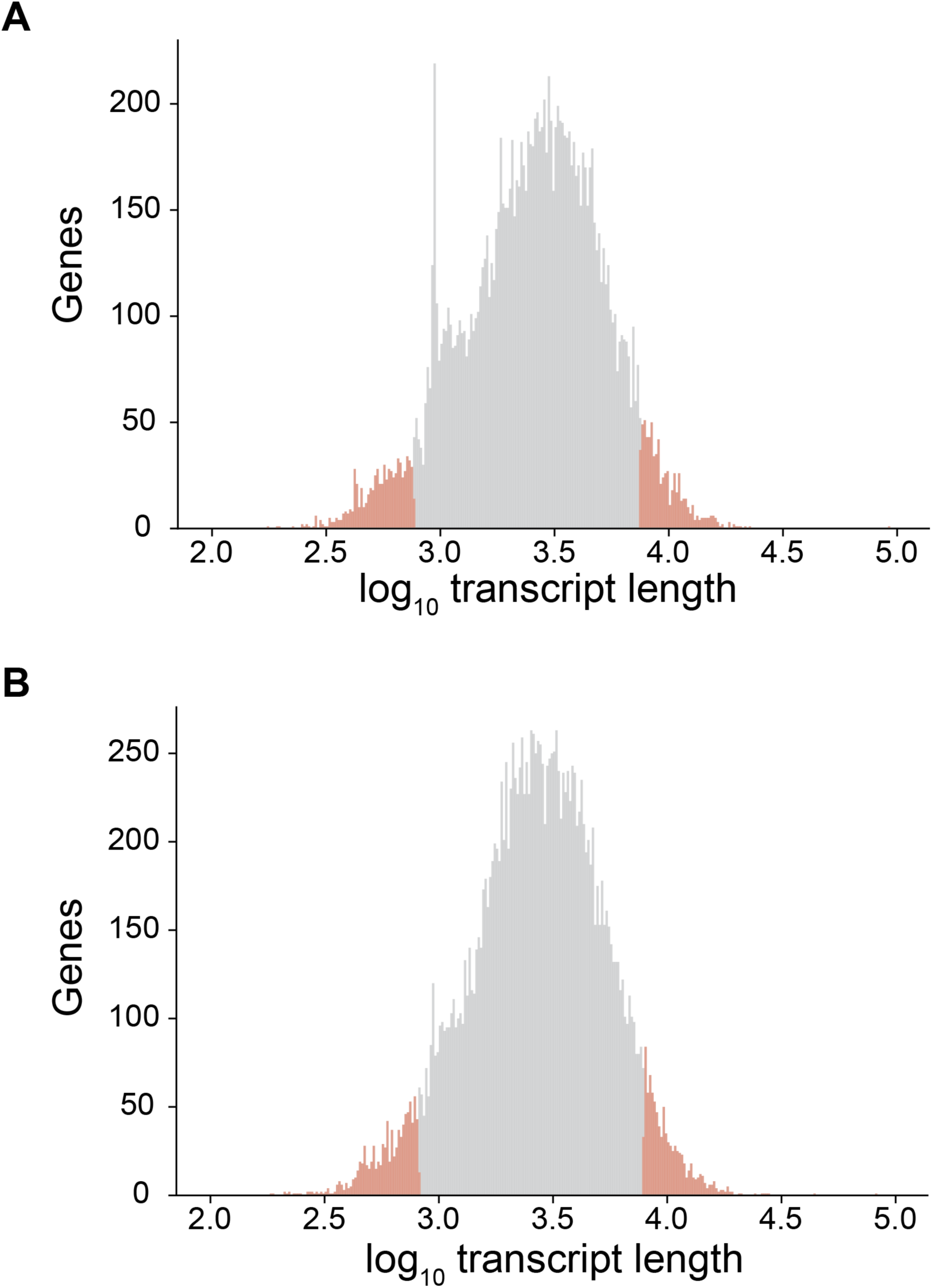
Distribution of median transcript length. (**A**) For mouse protein-coding genes. (**B**) For human protein-coding genes. Red indicates the genes with the 5% shortest and 5% longest transcripts.

**Fig. S18.**
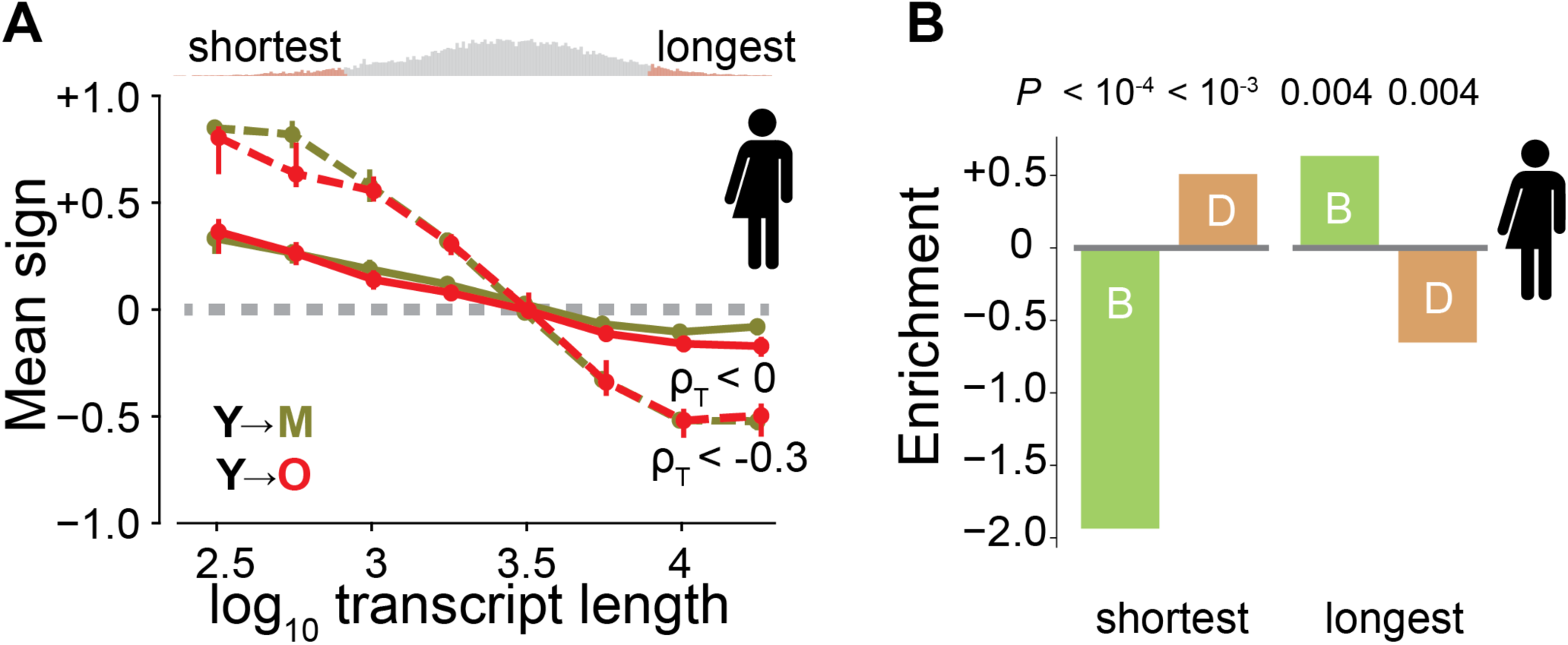
Human changes with transcript length. (**A**) Direction of age-dependent change of transcripts, analogous to Fig. 4A, but for humans. Additionally, the dotted curve shows samples with strong imbalance (ρ_IB_ < −0.3). The shortest and longest genes get most affected by transcriptome imbalance. (**B**) Fold enrichment for beneficial (B, green) and deleterious (D, orange) genes among the genes with the 5% shortest and 5% longest median transcript lengths in humans. Negative enrichment indicates depletion.

**Fig. S19.**
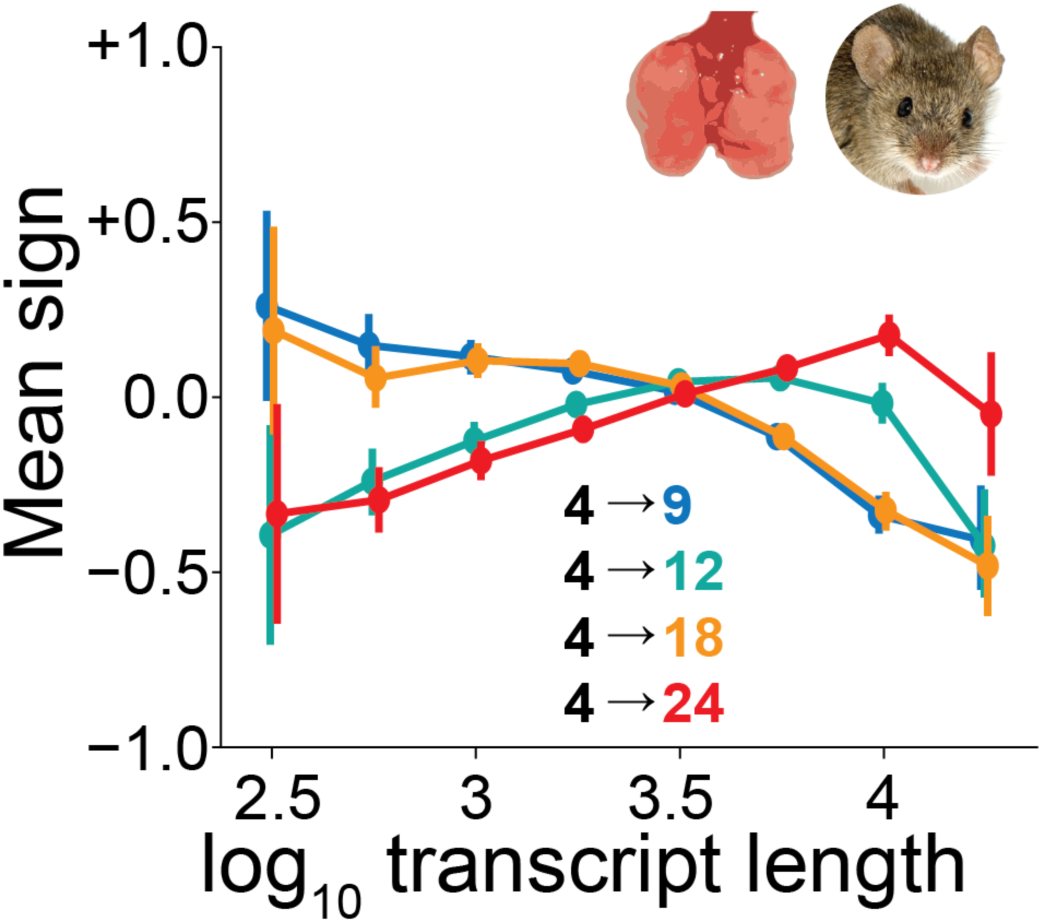
Changes with transcript length in uninfected lung of mice. An average sign of +1 would indicate that all genes are upregulated, whereas an average sign of −1 would indicate that all are downregulated.

**Fig. S20.**
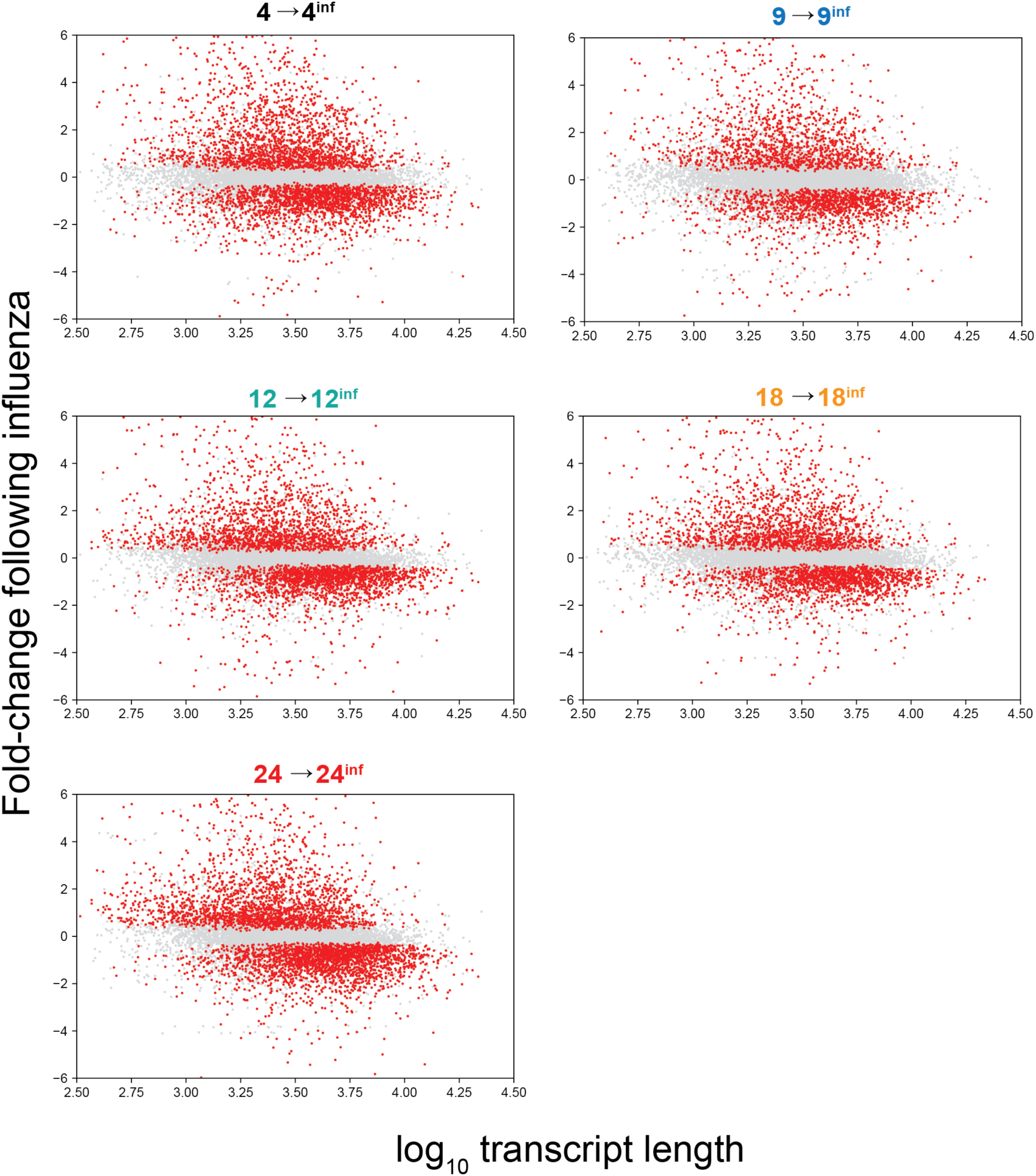
Transcript length dependency following influenza. Fold-changes observed in lung after influenza relative to lung without influenza exposure for individual genes. Graphs show mice of different ages. Red dots indicate differential expression at false discovery rate of <0.05.

**Table S1.**

Importance of individual contributing features.

**Table S2.**

Importance of individual contributing features in human GTEx.

**Table S3.**

Annotations enriched among human genes with short transcripts.

**Table S4.**

Annotations enriched among human genes with long transcripts.

**Table S5.**

Annotations enriched among mouse genes with short transcripts.

**Table S6.**

Annotations enriched among mouse genes with long transcripts.

**Table S7.**

Mouse genes correlating with transcriptome imbalance.

**Table S8.**

Correlation between transcript length and fold-changes of mouse studies in EBI-GXA.

**Table S9.**

Correlation between transcript length and fold-changes of human studies in EBI-GXA.

